# Expression of GRINA Correlates with Prognosis in Human Cancers: A Pan-cancer Analysis

**DOI:** 10.1101/2021.05.13.444089

**Authors:** S. M. Riazul Islam, Subbroto Kumar Saha, Shaker El-Sappagh, Faisal Tariq, Joydeep Das, Muhammad Afzal, Ssang-Goo Cho

## Abstract

*GRINA* is an emerging target for cancer therapy. However, the role of *GRINA* expression and its correlation with cancer patient survival has not been comprehensively studied. Here, we found that mRNA and protein expression of GRINA was upregulated in breast, colon, gastric, and prostate cancers and negatively correlated with patient survival. Also, the upregulation of GRINA expression is associated with hypomethylation of its promoter region. Our GRINA-miRNAs network analysis revealed potential regulatory miRNAs regulating the GRINA expression and its downstream pathways. Next, functional enrichment and pathway analysis of genes commonly co-express with *GRINA* in breast, colon, gastric, and prostate cancers revealed *GRINA* regulatory pathways. Concurrently, our upstream regulator analysis revealed possible kinases, transcription factors, and proteins that may potentially regulate *GRINA*. Overall, this study demonstrates the prognostic significance of *GRINA* expression and identifies potential regulatory mechanisms, which might have significant implications in targeted therapies for human cancers.

## 1. Introduction

Targeted therapy is an emerging paradigm in molecular medicine for next-generation cancer treatment. Recently, targeting the N-methyl-D-aspartate (NMDA) receptor expressed on the surfaces of cancer cells has emerged as a therapeutic approach [1]. NMDA receptors regulate the mammalian target of rapamycin (mTOR), a kinase involved in signaling that is a therapeutic target for many types of cancers [2–4]. Triggering the inappropriate expression of NMDA receptors on cancer cell lines thus represents a possible therapeutic avenue to control dysregulated cancer cell growth, division, and invasiveness [5, 6]. For example, NMDA receptors have been closely associated with tumor progression [7]. Inactivation of NMDA receptors in breast and lung cancers can potentially trigger apoptosis of tumor cells, and the NMDA receptor is a prospective therapeutic marker for ovarian cancer [8]. Also, NMDA2B phosphorylation can inhibit epileptic seizures triggered by brain tumors [9]. An essential constituent of NMDA heteromers in the NR2A subunit, the loss of which in gastric cancers can cause cell cycle arrest and prevent the proliferation of MKN45 [10].

One of the most common receptors belonging to the NMDA receptor family is GRINA, also known as transmembrane BAX inhibitor motif-containing (TMBIM3). GRINA, like other TMBIM members, is a regulator of cell death. All TMBIM family members inhibit different aspects of apoptosis, including ER modulation, calcium homeostasis, and ER stress signaling [11]. ER stress has also been shown to trigger TMBIM3/*GRINA* expression in defense against ER stress-triggered apoptosis [12]. GRINA also plays significant roles in cell proliferation, invasion, and migration [13], is strongly expressed in the central nervous system [14], and is overexpressed in breast cancer tissue [15]. Besides, *GRINA* mRNA is significantly upregulated in the prefrontal cortex of human subjects with a depressive disorder [16]. These findings suggest that GRINA may play a significant role in the progression of multiple cancers.

The existence of the volume of research on TMBIM members and their dysregulation in various cancer types [17–19] implies that the *GRINA* gene could be useful in finding innovative approaches for specific cancer therapies. *GRINA* mRNA levels as diagnostic markers in cancers can be considered an emerging area of research [11, 13, 14]. However, to the best of our knowledge, this gene has not been studied from a data mining perspective yet. In this study, we use multi-omics analysis to systematically evaluate the biomarker importance and prognostic significance of *GRINA* for multiple cancers. *GRINA* expression patterns and clinical outcomes of certain cancers were compared using expression, function, and patient survival datasets accessible online. We also investigated several regulatory mechanisms that may underlie *GRINA* function in our cancers of interest by probing for miRNA-mRNA interactions, co-expressed proteins, pathway activity, and interactions between kinases and transcription factors. This bioinformatics analysis ultimately demonstrated that *GRINA* expression could be used as a biomarker for determining prognoses for patients with certain types of cancers.

## 2. Experimental Section

### 2.1. GRINA mRNA and Protein Expression, Promoter Methylation, and Copy Number Alterations (CNAs) Analysis

The Oncomine platform (https://www.oncomine.org/resource/login.html) was used to analyze and visualize the *GRINA* mRNA expression [20–23]. Default threshold parameters were selected, consisting of *p*-value = 1E-4, fold-change = 2, and gene ranking in the top 10%. Statistical analyses were performed using an unpaired *t*-test, and *p* < 0.05 was considered significant.

*GRINA* mRNA expression in various types of cancer and normal tissues was examined using TCGA level 3 RNA-seq datasets via UCSC Xena (https://xenabrowser.net/heatmap/) [24]. Statistical analyses were performed using a Welch’s t-test, and *p* < 0.05 was considered significant.

*GRINA* mRNA expression in subclasses of multiple cancers, including breast, colon, gastric, and prostate cancers, was also determined using TCGA level 3 RNA-seq datasets via ULCAN (http://ualcan.path.uab.edu/index.html) [25]. Methylation values of *GRINA* gene promoter in cancers, such as breast, colon, gastric, and prostate cancer, were examined using TCGA datasets via ULCAN. Statistical analyses were performed using a student’s t-test, and *p* < 0.05 was considered significant.

*GRINA* mRNA expression in various cancer cell lines was analyzed by RT-PCR. For RT-PCR analysis, total RNA was extracted from used cell lines using the Easy-Blue RNA Extraction kit (iNtRON Biotechnology, Seongnam-si, Gyeonggi-do, Korea). According to the manufacturer’s instructions, total RNA (2 µg) was reverse transcribed into cDNA using a cDNA synthesis kit (Promega, Madison, WI, USA). The RT-PCR was assessed using r-Taq plus Master Mix (Elpis Biotech, Daejeon, Korea), and the PCR products were analyzed by ∼1.5% agarose gel electrophoresis. The bands were separated in agarose gels containing ethidium bromide (EtBr) and observed under UV light. The pictures were analyzed in Photoshop CS6 (Version 13.0.6 x64, San Jose, CA, USA), and the relative expression fold changes were measured in ImageJ [26]. The primers were used as Forward primer 5’-ATTCTCTGCATCTTCATCCGG-3’; Reverse primer 5’-AAACACATACTCTTCTGGGCTC-3’ for *GRINA* and Forward primer 5’-AATCCCATCACCATCTTCCAG-3’; Reverse primer 5’-CACGATACCAAAGTTGTCATGG-3’ for *GAPDH*.

GRINA protein expression in cancerous and normal tissues was examined using immunohistochemistry (IHC) staining data from the Human Protein Atlas (https://www.proteinatlas.org/) [27, 28]. For IHC staining, the HPA036980 antibody targeting GRINA was used. Copy number alterations (CNAs) of *GRINA* in various types of cancer were examined using TCGA datasets via cBioPortal (https://www.cbioportal.org/) [29, 30].

### 2.2. Prognosis Analysis Using R2: Kaplan Meier Scanner (Pro) and SurvExpress

The prognostic relationship between *GRINA* levels and cancer patient survival was investigated using the R2: Kaplan Meier Scanner web (https://hgserver1.amc.nl/cgi-bin/r2/main.cgi) [31] and using SurvExpress web (http://bioinformatica.mty.itesm.mx:8080/Biomatec/SurvivaX.jsp) [32]. Survival curve analysis was conducted using a threshold log-rank test using scan modus of expression value. Statistical significance was indicated by *p* < 0.05.

### 2.3. Construction of the GRINA-miRNA-mRNA Regulatory Network

miRNAs targeting *GRINA* mRNA were predicted and retrieved from miRSystem (http://mirsystem.cgm.ntu.edu.tw/index.php) [33] and starBase v3.0 (http://starbase.sysu.edu.cn/)[34]. Retrieved miRNAs were visualized as a Venn diagram using Venny v1.2 (https://bioinfogp.cnb.csic.es/tools/venny/) [35]. The *GRINA*-miRNA-mRNA regulatory network was constructed using starBase v3.0 to identify potential *GRINA*-binding miRNAs. Expression levels of these miRNAs in breast cancer were also determined using starBase v3.0 (http://starbase.sysu.edu.cn/). A Student’s t-test was performed to generate a *p*-value, which indicates the significance of an observation. *p* < 0.05 was considered statistically significant. The binding site of GRINA 3’UTR and miRNAs were derived from Starbase v3.0 web.

### 2.4. Gene Ontology (GO) and Pathway Analysis of GRINA Co-expressed Genes

Genes positively and negatively co-expressed with *GRINA* in selected cancers were retrieved from the TCGA data using R2: Genomics Analysis and Visualization Platform (https://hgserver1.amc.nl/cgi-bin/r2/main.cgi?&species=hs) [31]. Positively and negatively co-expressed genes were considered as those with r value −0.30 > and > 0.30 and P ≤ 0.01. Genes showing co-expression with *GRINA* in breast, colon, stomach, and prostate cancers were assembled into a Venn diagram via InteractiVenn (http://www.interactivenn.net/index.html) to identify the common positively and negatively co-expressed genes of GRINA [36]. The common genes co-expressed with *GRINA* in breast, colon, stomach, and prostate cancers were then used to analyze gene ontology and pathway enrichment using Enrichr (https://amp.pharm.mssm.edu/Enrichr/) [37, 38]. The bar graphs retrieved from Enrichr web were ranked by *p*-value from several databases, including the Kyoto Encyclopedia of Genes and Genomes (KEGG)-2019, Reactome pathway-2016, and GO (biological process, molecular function, and cellular component)-2018.

### 2.5. Transcription Factors and Protein Kinases Associated with Genes Co-expressed with GRINA

Upstream regulators and protein kinases associated with genes co-expressed with *GRINA* were identified by submitting the list of co-expressed genes to Expression2Kinases (X2K) web interface (https://amp.pharm.mssm.edu/X2K/) [39, 40], which identifies enriched transcription factors (TFs) upstream of the co-expressed genes using the ChEA database. Genes2Networks (G2N) module of X2K connected TFs with protein-protein interaction to identify transcriptional complexes related to these gene signatures. Protein kinases responsible for TF complex formation and regulation were recognized through the Kinase Enrichment Analysis (KEA) module of X2K. The top 10 most enriched TFs and kinases were ranked based on a combined P-value and z-score value.

To have a quick review of the aforementioned online tools, we have presented a schematic diagram that summarizes their functions in Figure 1.

**Figure 1.**
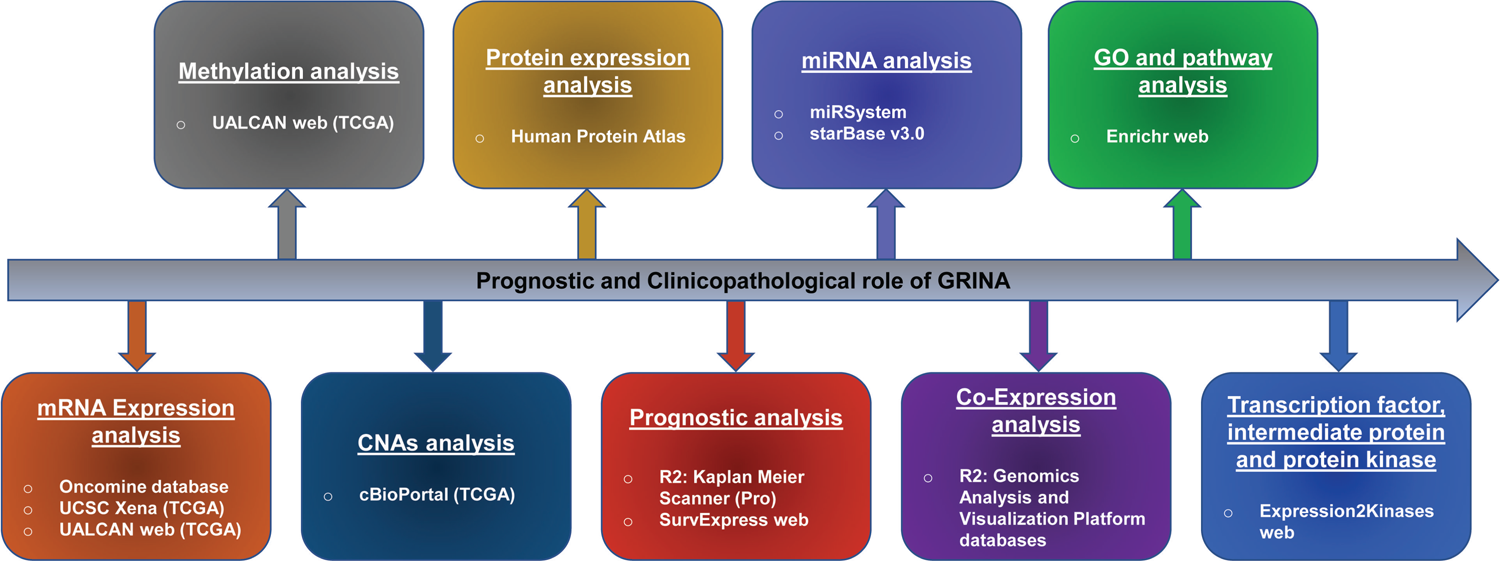
A schematic diagram summarizing selected online tools used in this study.

## 3. Result

### 3.1. GRINA mRNA Expression in Various Cancers

To examine *GRINA* mRNA expression levels in select normal and cancer types tissues, we utilized online analytical tools and databases, including Oncomine and TCGA. A significant number of analyses performed using the Oncomine platform revealed high *GRINA* expression in breast, colon, gastric, lymphoma, melanoma, ovarian, and prostate cancers (Figure 2a; Supplementary Table 1). The Oncomine platform also indicated the downregulation of *GRINA* in oesophageal, head-and-neck, and lung cancers (Figure 2a; Supplementary Table 1). To confirm the relative levels of *GRINA* expression in normal and cancerous tissues, we further analyzed expression data using the TCGA data-driven UCSC Xena online tool, which yielded similar results to those from Oncomine-based analysis (Figure 2b; Supplementary Table 2). We further experimentally confirmed the expression pattern of *GRINA* in various cancer cell lines using RT-PCR analysis (Figure 2c). According to the experimental outcomes, the mRNA expression of *GRINA* was upregulated in the breast (MCF7 and MDA-MB231 cell lines), colon (HCT116 and HT-29 cell lines), blood (K562 cell line), and ovarian (SKOV3 and A2780 cell lines). At the same time, it was comparatively downregulated in the esophagus (SEG-1 cell line), liver (HepG2 and SNU475 cell lines), and lung (A549 cell line) cancer. In summary, analysis with Oncomine and TCGA databases commonly showed *GRINA* overexpression in multiple cancers. Therefore, based on high *GRINA* expression levels, we selected breast, colon, gastric, and prostate cancers for further systematic analysis of *GRINA* expression and its clinical significance. It is worth noting that we used Oncomine and TCGA databases to present the expression status of *GRINA* in various cancers. The primary purpose of this expression checking was to find the four most common cancers in terms of high *GRINA* expression. It can be noted that our purpose of the multi-omics analysis is not to see the *GRINA* expression status individually. Instead, we used multi-omics analysis to show consistencies between *GRINA* expression and other genetic statuses such as methylation of *GRINA* promoters, mutation and CNA, and immunostaining results, as will be analyzed subsequently.

**Figure 2.**
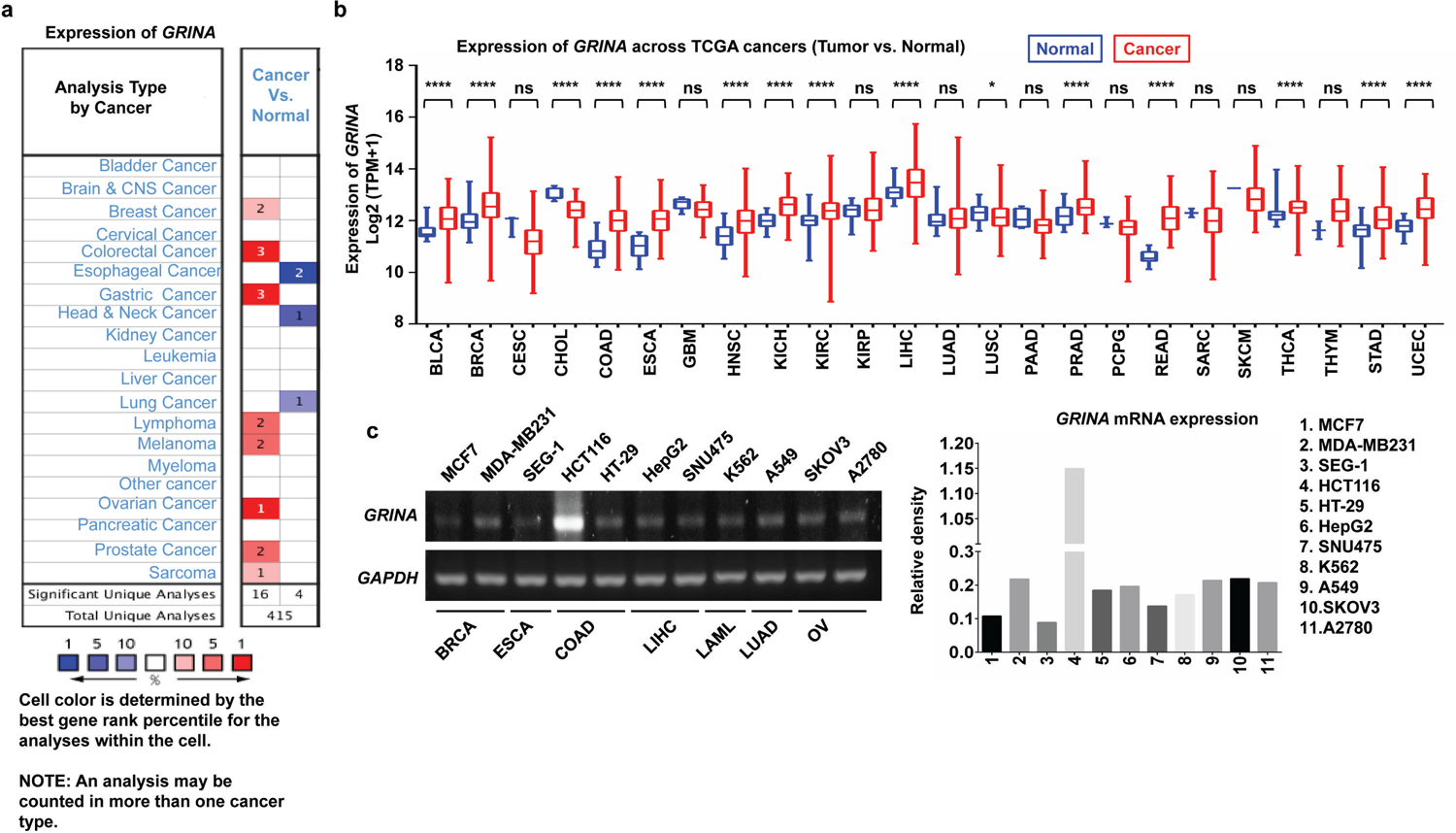
*GRINA* (Glutamate Ionotropic Receptor NMDA Type Subunit Associated Protein 1) mRNA levels were analyzed in different types of cancer. (**a**) The mRNA expression of *GRINA* in various human cancers. The figure was retrieved from the Oncomine database. The cell number represents the dataset number that meets all the thresholds with the color blue for under-expression and color red for over-expression. Cell color is determined by the best gene rank percentile for the analyses within the cell. (**b**) The mRNA expression of *GRINA* in various human cancers. The figure was retrieved from the TCGA database using UCSC Xena. *p*-value < 0.05 represents statistically significant. \**p* < 0.05; ** *p* < 0.01; \*\*\**p* < 0.001; \*\*\*\**p* < 0.0001. (**c**) mRNA expression of *GRINA* was analyzed by RT-PCR using various cancer cell lines including breast (MCF7 and MDA-MB231), esophagus (SEG-1), colon (HCT116 and HT-29), liver (HepG2 and SNU475), blood (K562), lung (A549), and ovarian (SKOV3 and A2780), cancers. *GAPDH* was used as a loading control. The expression intensity of *GRINA* mRNA was analyzed by ImageJ software. The expression bands were cropped from the full gel with no apparent modification. The image of full gels is supplemented in supplementary information files under Supplementary Figure 1.

### 3.2. GRINA Expression Patterns and Patient Survival of Breast Cancer

To investigate *GRINA* expression in breast cancer and corresponding normal tissue samples, we evaluated Oncomine and TCGA datasets. Relative expression levels of *GRINA* in 21 lobular breast cancer (LBC) and 38 invasive ductal breast cancer (IDBC) tissues were analyzed in the Zhao dataset. Compared to histologically normal breast tissue (*n* = 3) derived from IDBC patients used in the Zhao dataset, *GRINA* is significantly overexpressed in both LBC and IDBC tissues (Figure 3a-b). In addition, *GRINA* mRNA levels are significantly (*p* < 1.63E-12) higher in invasive breast carcinoma (BRIC) tissue than normal tissue (Figure 3c) based on the analysis of 114 normal breast tissue samples and 1097 breast cancer tissue samples from the TCGA-driven ULCAN dataset. To analyze GRINA expression and patient subclass associations, we determined *GRINA* levels in normal, luminal, human epidermal growth factor receptor 2 positive (HER2+), and triple-negative breast cancer positive (TNBC+) groups, which were derived from 833 patient samples in the TCGA dataset using ULCAN. *GRINA* mRNA levels are significantly higher in luminal (*p* = 1.63E−12), HER2+ methylation level of *GRINA* gene promoters in breast cancer samples is significantly lower compared to that in normal breast samples (Figure 3e). A lack of methylation of *GRINA* promoters in breast cancer may thus cause *GRINA* upregulation of this gene. We next analyzed mutation and copy number alteration (CNA) of *GRINA* in breast cancer using cBioPortal. Alterations of the *GRINA* gene, including amplification and deep deletion, were found in 32% of 1088 cases (Figure 3f). *GRINA* mRNA levels were high in approximately 50% of altered samples, suggesting that *GRINA* expression in breast cancer is positively correlated with CNA status. Such amplification likely underlies increases in *GRINA* expression.

**Figure 3.**
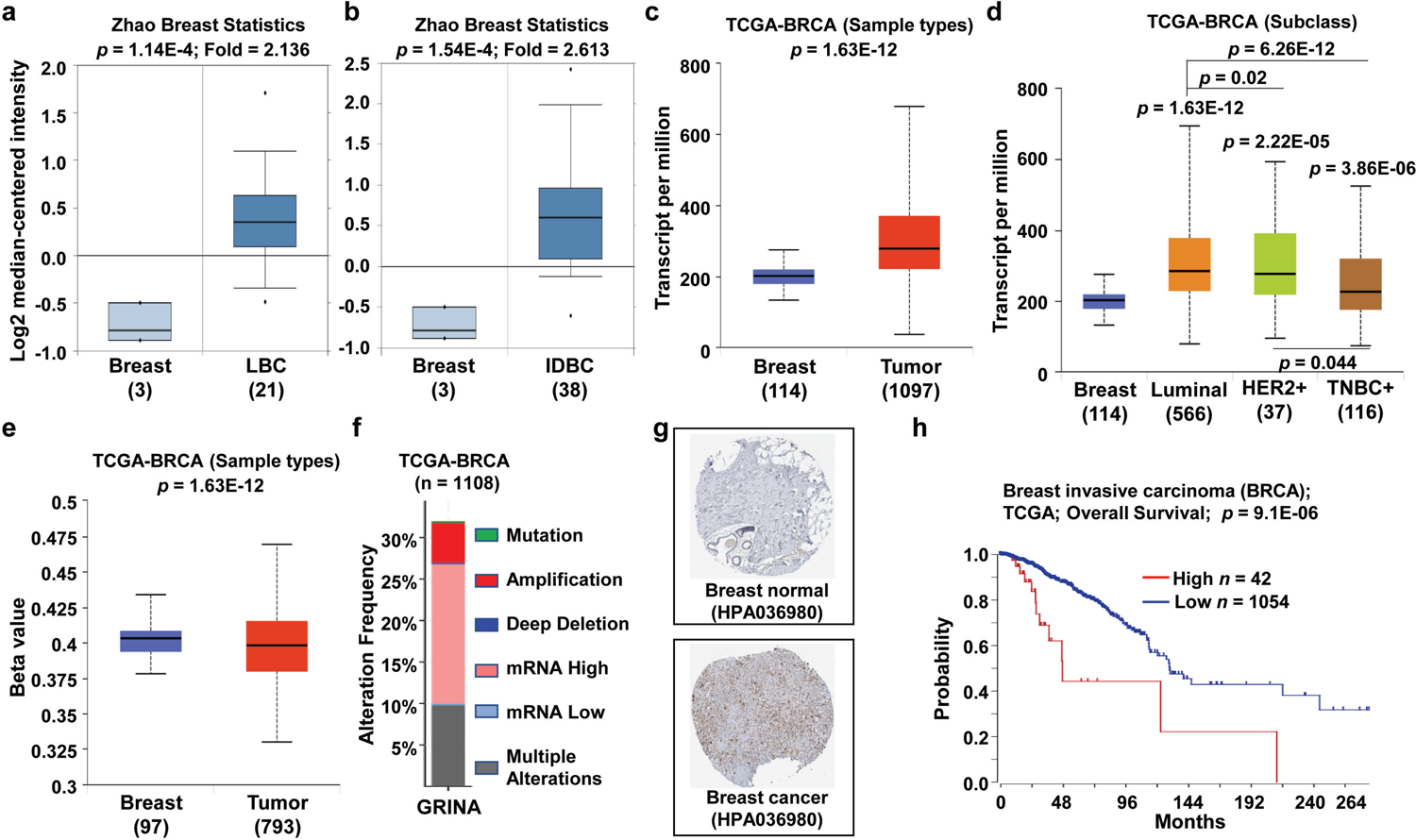
Expression of *GRINA* and patient survival were analysed in normal and cancerous breast tissue. (**a and b**) Fold-changes in *GRINA* levels in breast cancer, shown as box plots comparing *GRINA* expression in normal (*n* = 3, left plot) vs. lobular breast carcinoma tissue (*n* = 21, right plot) (**a**) and in normal (*n* = 3, left plot) vs. invasive ductal breast carcinoma tissue (*n* = 38, right plot) (**b**). Data derived from the Oncomine database. (**c**) *GRINA* mRNA levels were analyzed in breast cancer (n = 1097) and the normal (n = 114) tissues based on data from The Cancer Genome Atlas (TCGA) database using ULCAN web. (**d**) *GRINA* levels in BRCA patients (subclass) based on the TCGA database using ULCAN web. (**e**) Methylation levels in *GRINA* promoters were analysed in breast cancer based on The Cancer Genome Atlas (TCGA) data and ULCAN. (**f**) Mutation and copy number alteration frequencies of *GRINA* were derived from cBioPortal using TCGA-BRCA data. (**g**) GRINA protein expression data from immunohistochemistry staining in normal and breast carcinomas was derived from the Human Protein Atlas. (**h**) Survival curves comparing breast cancer patients with high (red) and low (blue) *GRINA* levels based on the R2: Kaplan Meier Scanner (Pro) database. Survival curve analysis was conducted using a threshold logrank test using scan modus of expression value. *p*-value < 0.05 represents statistically significant.

Protein expression patterns of GRINA in breast cancer were examined using immunohistochemical (IHC) staining and the Human Protein Atlas. These results confirm the overexpression of *GRINA* at the protein levels in breast cancer samples relative to normal breast tissue (Figure 3g). In addition, we investigated the relationship between *GRINA* expression and clinical prognosis using the R2: Kaplan Meier Scanner. Patients with high levels of *GRINA* (*n* = 42) had significantly (*p* = 9.1E-06) lower overall survival compared to patients with lower *GRINA* levels (*n* = 1054) (Figure 3h). *GRINA* levels are thus significantly augmented in breast cancer cells and positively correlated with poor patient prognosis.

### 3.3. GRINA Expression Patterns and Patient Survival of Colon Cancer

We next examined *GRINA* characteristics in colon cancer (Figure 2a-d). Upregulation patterns of *GRINA* in colon cancer have previously been reported [11, 41]; however, the correlation between *GRINA* expression and patient prognosis has not been systematically analyzed. We thus evaluated *GRINA* expression in microarray datasets for colon cancer and normal tissues using Oncomine. In addition, relative expression levels of *GRINA* in colon carcinomas (CC) and 65 rectal adenocarcinomas were analyzed using the Skrzypczak and Gaedcke datasets. Compared to normal colon tissue, *GRINA* was significantly upregulated in both cases (Figures 4a and b). We also analyzed *GRINA* levels in 41 colon normal tissue samples and 286 colon cancer tissue samples. As observed for breast cancer samples, *GRINA* expression was significantly (*p* < 1E−12) augmented in colon adenocarcinomas (COAD) compared to normal tissues (Figure 4c).

**Figure 4.**
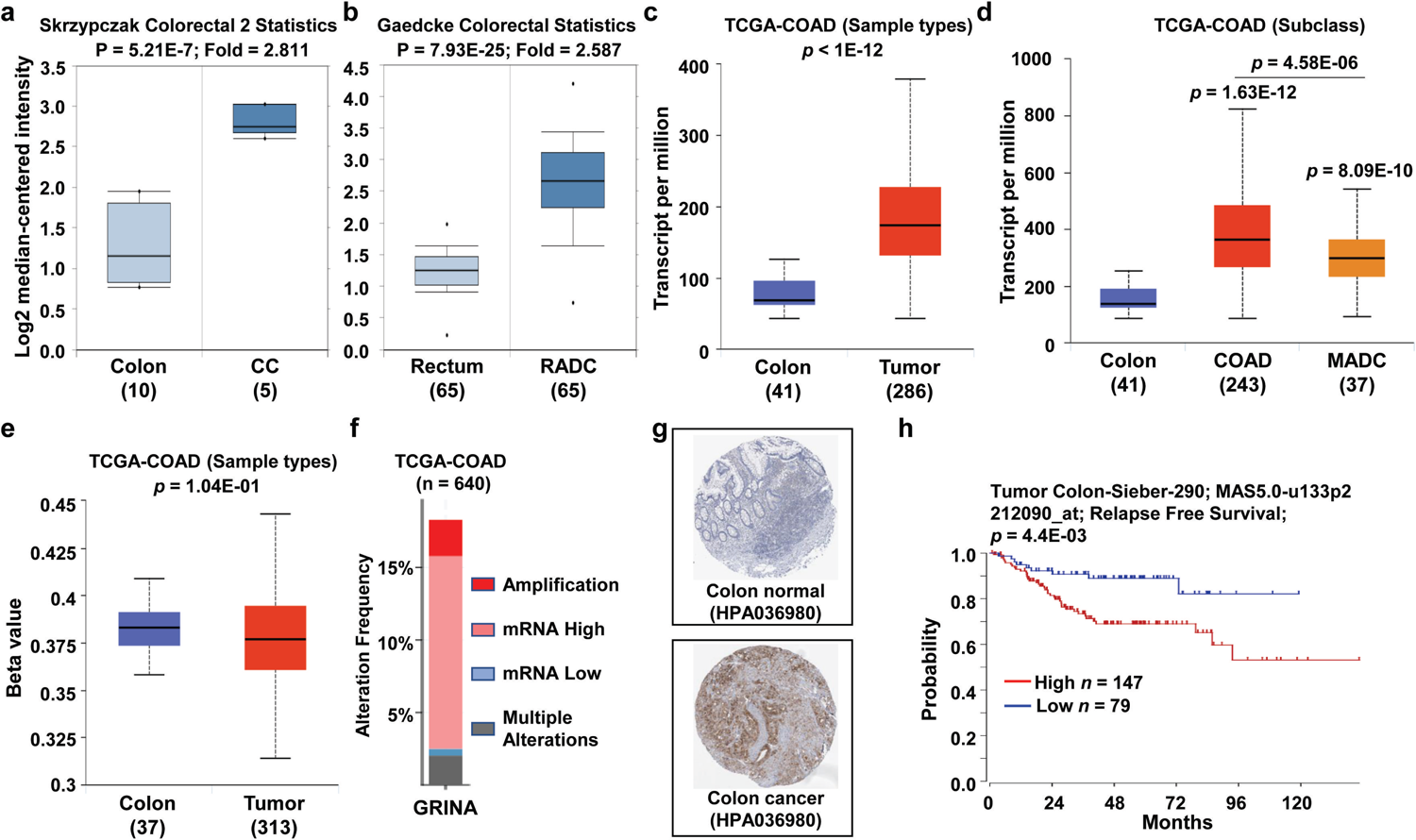
*GRINA* expression patterns and patient survival analysis were performed for colorectal cancers. (**a and b**) Fold-changes in *GRINA* in colorectal cancers were identified by our analyses, shown as a box plot comparing *GRINA* levels in normal (*n* = 10, left plot) vs. colon carcinoma tissues (*n* = 5, right plot) (**a**) and normal (*n* = 65, left plot) vs. rectal adenocarcinoma tissue (*n* = 65, right plot) (**b**) based on data from the Oncomine database. (**c**) *GRINA* mRNA levels in colon adenocarcinoma were obtained from The Cancer Genome Atlas (TCGA) database through ULCAN. (**d**) *GRINA* expression in COAD patients (subclass) obtained from the TCGA database. (**e**) Methylation of *GRINA* gene promoters was analyzed in colon cancer based on data from The Cancer Genome Atlas (TCGA) database obtained via ULCAN. (**f**) Copy number alteration frequencies of the *GRINA* gene were derived from the cBioPortal web using TCGA-COAD data. (**g**) Protein expression of GRINA, as demonstrated by immunohistochemistry staining in normal tissue and colon carcinomas obtained from the Human Protein Atlas. (**h**) Survival curves for colon cancer patients with high (red) and low (blue) GRINA levels in colon cancer based on the R2: Kaplan Meier Scanner (Pro) database. Survival curve analysis was conducted using a threshold logrank test using scan modus of expression value. *p*-value < 0.05 represents statistically significant.

We also analyzed 321 patient samples in the TCGA dataset to examine the relationship between *GRINA* expression and colon cancer patient subclass. GRINA mRNA levels were significantly elevated in COAD (*p* = 1.63E−-12) and mucinous-adenocarcinoma (MADC) (*p* = 8.09E−10) subclasses (Figure 4d), which is supported by the hypomethylation of the GRINA gene promoter in colon cancer specimens. However, the difference between normal and cancer is not statistically significant (Figure 4e). In addition, mutation and CNA analysis of *GRINA* in colon cancer samples using cBioPortal revealed that *GRINA* levels positively correlated with CNA status, as 85 out of 117 colon cancer cases with altered samples showed high *GRINA* mRNA levels (Figure 4f). While amplification occurred in a handful of cases, no specific mutation appeared to cause *GRINA* alterations.

We also examined GRINA protein expression in colon cancer specimens using IHC overexpression was visible across the entire colon cancer tissue sample compared to normal tissue. We then analyzed colon cancer patients’ clinical outcomes to understand the relationship between prognosis and *GRINA* expression. The R2: Kaplan Meier Scanner was applied to data from 147 patients with high *GRINA* expression and 79 patients with low *GRINA* expression. Patients with high *GRINA* levels experienced poorer relapse-free survival (Figure 4h). Thus, these results suggest that high GRINA expression due to copy number alterations are common in colon cancer tissue and positively correlates with low patient survival.

### 3.4. GRINA Expression Patterns and Patient Survival in Gastric Cancer

We also examined *GRINA* levels in gastric carcinoma (GC) in which *GRINA* overexpression has previously been observed [13, 42, 43]. We thus performed a systematic analysis to evaluate the association between *GRINA* expression and clinical outcomes of gastric cancer patients. Relative *GRINA* mRNA levels in 26 Gastric Intestinal Type Adenocarcinoma (GITA) and 4 Gastric Mixed Adenocarcinoma (GMA) were analyzed based on the DErrico datasets using the Oncomine platform. *GRINA* levels are significantly altered in both types of gastric cancers compared to normal gastric tissue (Figure 5a and b). Analysis of *GRINA* expression in a larger TCGA dataset consisting of 34 normal stomach tissue samples and 415 gastric cancer tissue samples confirmed expression patterns described above, as *GRINA* was significantly (*p* < 1.63E−12) overexpressed in gastric cancer samples (Figure 5c). Next, TCGA was used to access data from 440 patients and determine whether *GRINA* mRNA levels were significantly elevated in each gastric cancer grade (Figure 5d), expression levels alone did not always indicate the cancer stage, as Grade 3 gastric cancers exhibited the lowest levels of *GRINA*.

**Figure 5.**
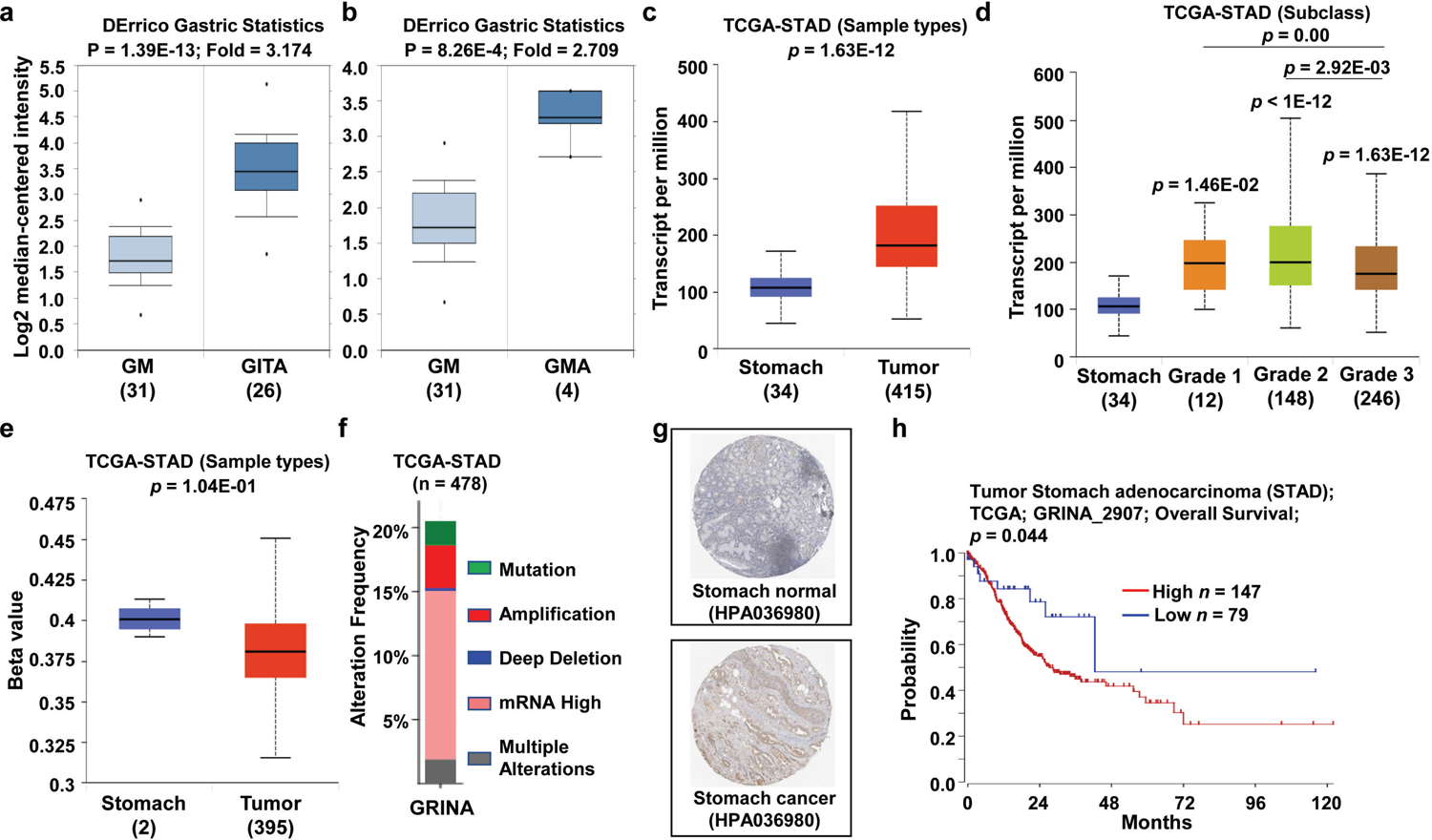
*GRINA* expression patterns and patient survival were analysed in gastric cancers. (**a and b**) Fold-changes of *GRINA* in gastric cancers is shown as box plots either comparing *GRINA* levels in normal (*n* = 31, left plot) and gastric intestinal-type adenocarcinoma tissues (*n* = 26, right plot) (**a**) or comparison normal (*n* = 31, left plot) and gastric mixed adenocarcinoma tissues (*n* = 4, right plot) (**b**) based on the Oncomine database. (**c**) *GRINA* mRNA levels in stomach carcinomas was obtained from The Cancer Genome Atlas (TCGA) database through ULCAN. (**d**) *GRINA* gene expression in STAD patients (subclass) was obtained from the TCGA database. (**e**) Methylation of *GRINA* gene promoters was analyzed in STAD tumors (red plot) and normal (blue plot) tissues based on TCGA database information and was generated using ULCAN. (**f**) Mutation and copy number alteration frequencies of the *GRINA* gene were derived from the cBioPortal web using TCGA-STAD data. (**g**) GRINA protein expression based on immunohistochemistry staining in normal and stomach carcinoma tissues was derived from the Human Protein Atlas. (**h**) Survival curves or stomach cancer patients were plotted with high (red) and low (blue) GRINA levels based on the R2: Kaplan Meier Scanner (Pro) database. Survival curve analysis was conducted using a threshold logrank test using scan modus of expression value. *p*-value < 0.05 represents statistically significant.

To evaluate the consistency in *GRINA* levels with *GRINA* promoter methylation, we compared the beta values of *GRINA* promoter methylation in normal gastric and cancerous gastric tissues. A large but non-significant difference in beta values between gastric cancer and normal samples was observed (Figure 5e), suggesting elevated *GRINA* mRNA levels in gastric cancer. Out of these altered cases, 64% (63 out of 98 altered cases) showed high *GRINA* mRNA levels (Figure 5f), indicating a positive association between CNA status and *GRINA* levels in gastric cancer. IHC results with gastric cancer specimens also decisively indicate increased *GRINA* din gastric cancer (Figure 5g). We next investigated whether *GRINA* expression likely relates to gastric cancer prognosis. Using the same number of patients as in the colon cancer analysis, R2: Kaplan Meier Scanner analysis of gastric cancer cases indicated an association between poor overall patient survival and higher *GRINA* mRNA levels (Figure 5h). Taken together, *GRINA* overexpression is expected in most gastric cancer cases and positively associates with poor clinical prognosis.

### 3.5. GRINA Expression Patterns and Patient Survival of Prostate Cancer

*GRINA* mRNA expression is known to be highly upregulated in prostate cancer tissues compared to normal tissues based on online tools and databases (Figure 6a-d). Our detailed analysis showed that elevated *GRINA* expression was evident in prostate carcinomas and prostatic intraepithelial neoplasias based on Oncomine (Figure 6a and b) using the Tomlins datasets. *GRINA* expression was also significantly (*p* = 4.82E-14) upregulated in prostate adenocarcinomas (PRAD) according to analysis using the TCGA database (Figure 6c) and considering 52 normal prostate tissue samples and 497 prostate cancer tissue samples. For prostate cancer, *GRINA* mRNA expression patterns can be translated into Gleason scores (GS), which range from 1 to 5, and describe whether prostate tissue biopsies resemble normal tissues (lower score) or abnormal tissues (higher score). As presented in Figure 6d, *GRINA* levels and the GS exhibited linear relationships, indicating that the most melancholy GRINA expression in prostate cancer represents GS 1. The highest *GRINA* levels represent GS 5, and *GRINA* levels corresponding to other GS values can be determined using a simple linear transformation.

**Figure 6.**
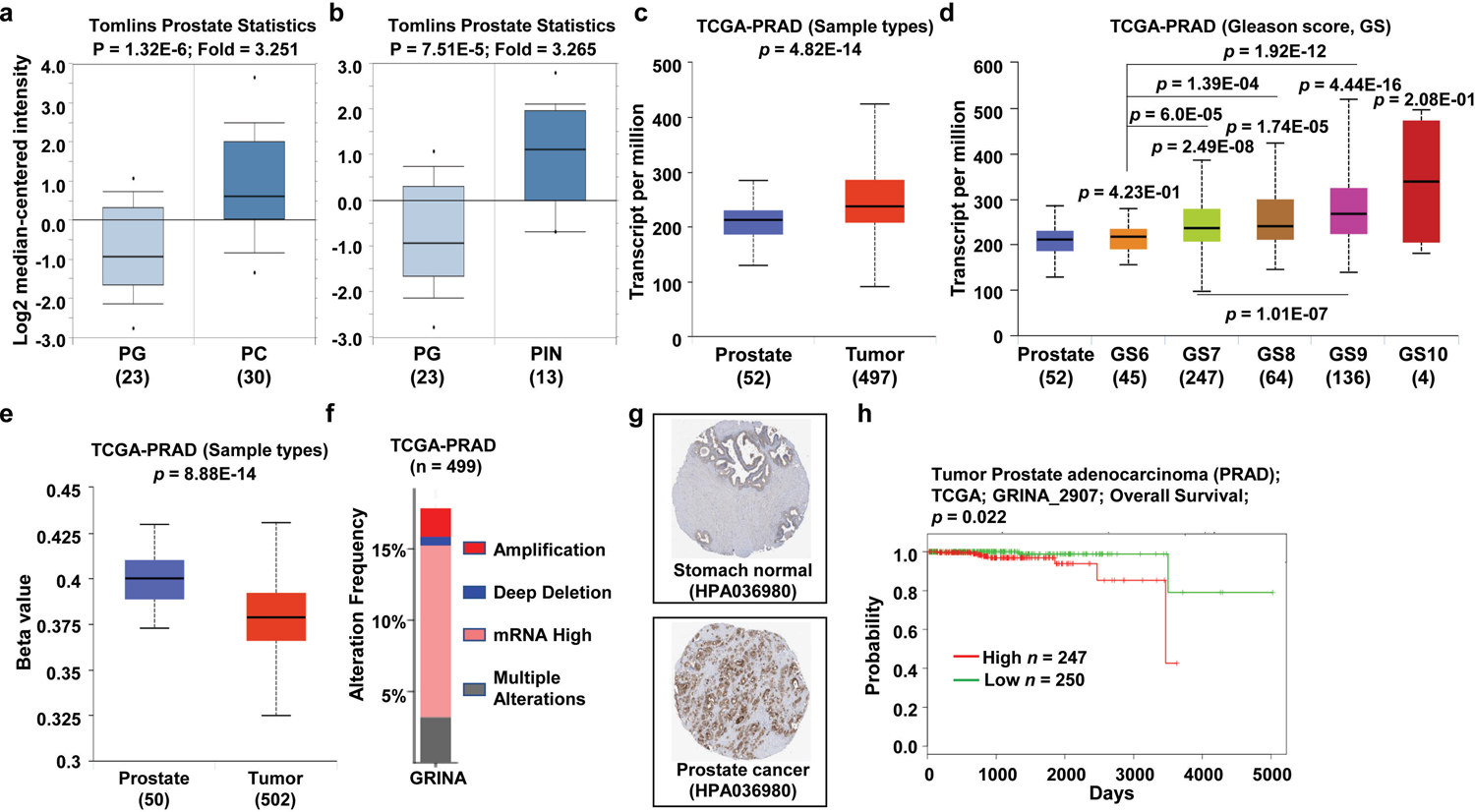
*GRINA* expression patterns and patient survival were analyzed for prostate cancer. (**a and b**) The fold-changes in *GRINA* levels in prostate cancers are shown as box plots comparing levels between normal (*n* = 23, left plot) and prostate carcinoma tissues (*n* = 30, right plot) (**a**) and between normal (*n* = 23, left plot) and prostatic intraepithelial neoplasia tissues (*n* = 13, right plot) (**b**) based on data derived from Oncomine. (**c**) *GRINA* mRNA levels was analyzed in prostate adenocarcinomas from The Cancer Genome Atlas (TCGA) database via ULCAN web. (**d**) *GRINA* expression levels was examined in PRAD patients (Gleason score) from the TCGA database. (**e**) Methylation of *GRINA* gene was analyzed in prostate cancer using The Cancer Genome Atlas (TCGA) database through the ULCAN web. (**f**) Copy number alteration frequency of the *GRINA* gene was derived from the cBioPortal web using TCGA-PRAD data. (**g**) Protein expression data for GRINA was obtained via immunohistochemistry staining in normal and prostate carcinoma tissues from the Human Protein Atlas. (**h**) Survival curves comparing patients with prostate cancer having high (red) and low (blue) GRINA levels, plotted using data from the R2: Kaplan Meier Scanner (Pro) database. Survival curve analysis was conducted using a threshold logrank test using scan modus of expression value. *p*-value < 0.05 represents statistically significant.

Methylation status analysis revealed that the *GRINA* gene promoter was significantly hypomethylated in prostate cancer tissues, confirming mRNA overexpression patterns of *GRINA* (Figure 6e). In addition, CNA status and *GRINA* expression levels in prostate cancer were positively correlated, with ∼ 67% (60 out of 89 altered cases) of altered samples showing high levels of *GRINA* mRNA (Figure 6f). Human Protein Atlas-assisted immunostaining confirmed the GRINA protein overexpression in prostate cancer samples compared to normal samples (Figure 6g). Finally, we investigated the prognostic significance of *GRINA* expression levels for prostate cancer. R2: Kaplan Meier Scanner survival analysis of data for a set of 247 patients with high *GRINA* levels and 250 patients with low *GRINA* levels revealed that prostate cancer patients with high *GRINA* levels showed significantly poorer overall survival compared to the low *GRINA* expression patients (Figure 6h). Overall, the available data demonstrated altered expression levels of *GRINA* in prostate cancer and their associated prognostic utility.

### 3.6. GRINA Expression Patterns and Patient Survival for Additional Cancer Types

Although the primary focus of this study was to understand *GRINA* expression levels and their clinical significance in the four cancers presented above, we also analyzed *GRINA* expression levels and their relevance to clinical outcomes in other cancers. Based on Oncomine analyses, *GRINA* expression was elevated in cancers such as lymphoma, melanoma, ovarian, and sarcoma cancers (Figure 2b). Using TCGA database-enabled ULCAN analysis, significant upregulation of *GRINA* was found in many additional cancers, including bladder, cholangial, oesophageal, head-and-neck, liver, rectum, and uterine cancers (Supplementary Figure 2). Our analysis did not identify any cancers in which *GRINA* was significantly (*p* < 0.05) downregulated. To understand the prognostic relevance of *GRINA* levels, analysis of survival rates and *GRINA* levels was performed using the SurvExpress platform based on TCGA data. This analysis showed that *GRINA* levels positively correlated with low patient survival in cancers such as the bladder, cholangial, and rectum cancers but not uterine cancer, as indicated by hazard ratios (Supplementary Figure 3). These findings suggested that *GRINA* levels are probably associated with biological mechanisms that promote aggressiveness to some extent in most cancers.

### 3.7. Analysis of Potential GRINA-miRNA-mRNA Regulatory Networks

After analyzing *GRINA* expression patterns and clinical relevance in various cancer types, we focused on a couple of underlying regulatory mechanisms of *GRINA* in our cancers of interest. Several studies [44, 45] suggest that microRNA (miRNA)-targeting genes can be utilized as prognostic predictors for individual cancer patients, conforming to the competing endogenous RNA (ceRNAs) hypothesis. We thus built and analyzed a potential *GRINA*-miRNA-mRNA regulatory network. Using the miRSystem, 41 miRNAs were found that potentially targeted *GRINA*, while starBase provided 54 such miRNAs (Figure 7a). A total of 18 of these miRNAs appeared on both platforms. It was chosen as candidate miRNAs for targeting *GRINA* mRNA to use with the starBase web to create a *GRINA*-miRNA-mRNA regulatory network (Figure 7b). As shown earlier, *GRINA* is upregulated in breast, colon, gastric, and prostate cancers. To conform with these mRNA expression patterns, expression levels of *GRINA*-targeting miRNAs should be lower in cancers compared to normal controls, according to the ceRNA model. We, therefore, examined expression levels of the 18 potential *GRINA*-targeting miRNAs in the aforementioned cancers. miR-411-5p, miR-654-5p, and miR-874-3p were marked as representative miRNAs because the expression levels of these miRNAs are significantly lower in cancerous tissues than other miRNAs. miRNA expression patterns of miR-411-5p, miR-654-5p, and miR-874-3p in the above cancers followed the ceRNAs mechanism, except for miR-411-5p in colon cancer and both miR-654-5p and miR-874-3p in gastric cancer (Figure 7c). In silico algorithms (Starbase v3.0) were also utilized to identify the binding sites of predicted miRNAs (miR-411, miR-654, and miR-874) that targeted the 3′ UTR region of the GRINA gene. The results showed that our predicted miRNAs were explicitly bound to the 3’UTR region of the GRINA gene, implying direct prevention of GRINA transcription via miR-411, miR-654, and miR-874 (Figure 7d (i-iii)). Our findings suggest that *GRINA* may have oncogenic roles in breast and prostate cancers via this miRNA-mRNA regulatory network. However, other unknown factors could overshadow the roles of the miRNA-mRNA network on the oncogenic behavior of *GRINA* in the colon and gastric cancers.

**Figure 7.**
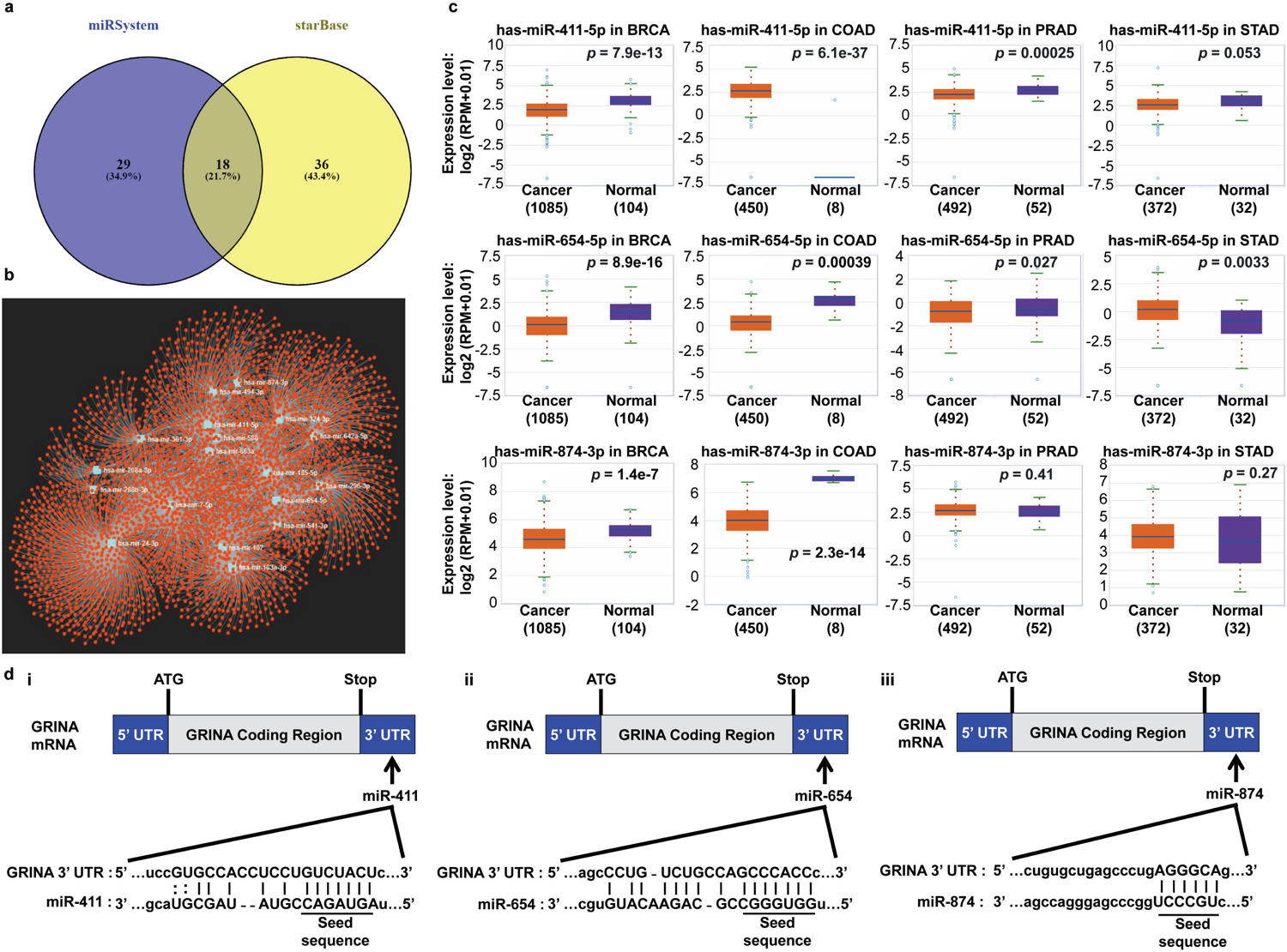
Construction of a GRINA-miRNA-mRNA network. (**a**) GRINA mRNA-targeting miRNAs were identified using miRSystem and starBase v3.0 databases. Common miRNAs were identified using list of miRNAs derived from miRSystem and Starbase v3.0 via Venn diagram. (**b**) A total of common 18 GRINA mRNA-targeting miRNAs were used to construct the GRINA-miRNA-mRNA network using the starBase v3.0 database. (**c**) Expression of miR-411-5p, miR-654-5p, and miR-874-3p was analysed in breast, colon, prostate, and stomach cancer tissues and normal tissue controls as determined using starBase v3.0. (d) Schematic of the in-silico analysis of the predicted binding sites showing that miR-411, miR-654, and miR-874 bind to the 3′ UTR region of GRINA mRNA. The binding site of GRINA 3’UTR and miRNAs were derived from Starbase v3.0 web. The predicted consequential pairing between the target region (position chr8; 145067249–145067271 of GRINA 3′ UTR) and the seed sequence of miR-411 is shown (i). The predicted consequential pairing between the target region (position chr8; 145067297–145067303 of GRINA 3′ UTR) and the seed sequence of miR-654 is shown (ii). The predicted consequential pairing between the target region (position chr8; 145067491–145067496 of GRINA 3′ UTR) and the seed sequence of miR-874 is shown (iii). *p*-value < 0.05 represents statistically significant.

### 3.8. Identification and Functional Enrichment Analysis of Genes Co-expressed with GRINA

To understand how *GRINA* functions in conjunction with other genes in signaling pathways in breast, colon, gastric, and prostate cancers, co-expression with *GRINA* was analyzed using TCGA data through the R2: Genomics Analysis and Visualization Platform. Eighty-three genes were upregulated with *GRINA* upregulation (“Positive Cluster”) (Figure 8a), while 21 genes were downregulated with *GRINA* upregulation (“Negative Cluster”) (Figure 8d). To understand the function of these genes, gene ontology (GO) functional annotation and pathway enrichment analysis were performed using the Enrichr platform. Three GO functional annotation categories for each cluster were obtained: biological process, cellular component, and molecular function. For the positive cluster, attachment of GPI (glycosylphosphatidylinositol) anchor to protein, GPI anchor binding, and mRNA cleavage and polyadenylation specificity factor complex were the most significantly represented GO terms in the biological process, cellular component, and molecular function, respectively (Supplementary Figure 4 a-c). Conversely, entirely different sets of ontologies were component is mutLapha complex, with several critical biological processes including DNA binding, siRNA binding, and helicase activity and molecular functions related to this cluster (Supplementary Figure 4 d-f).

**Figure 8.**
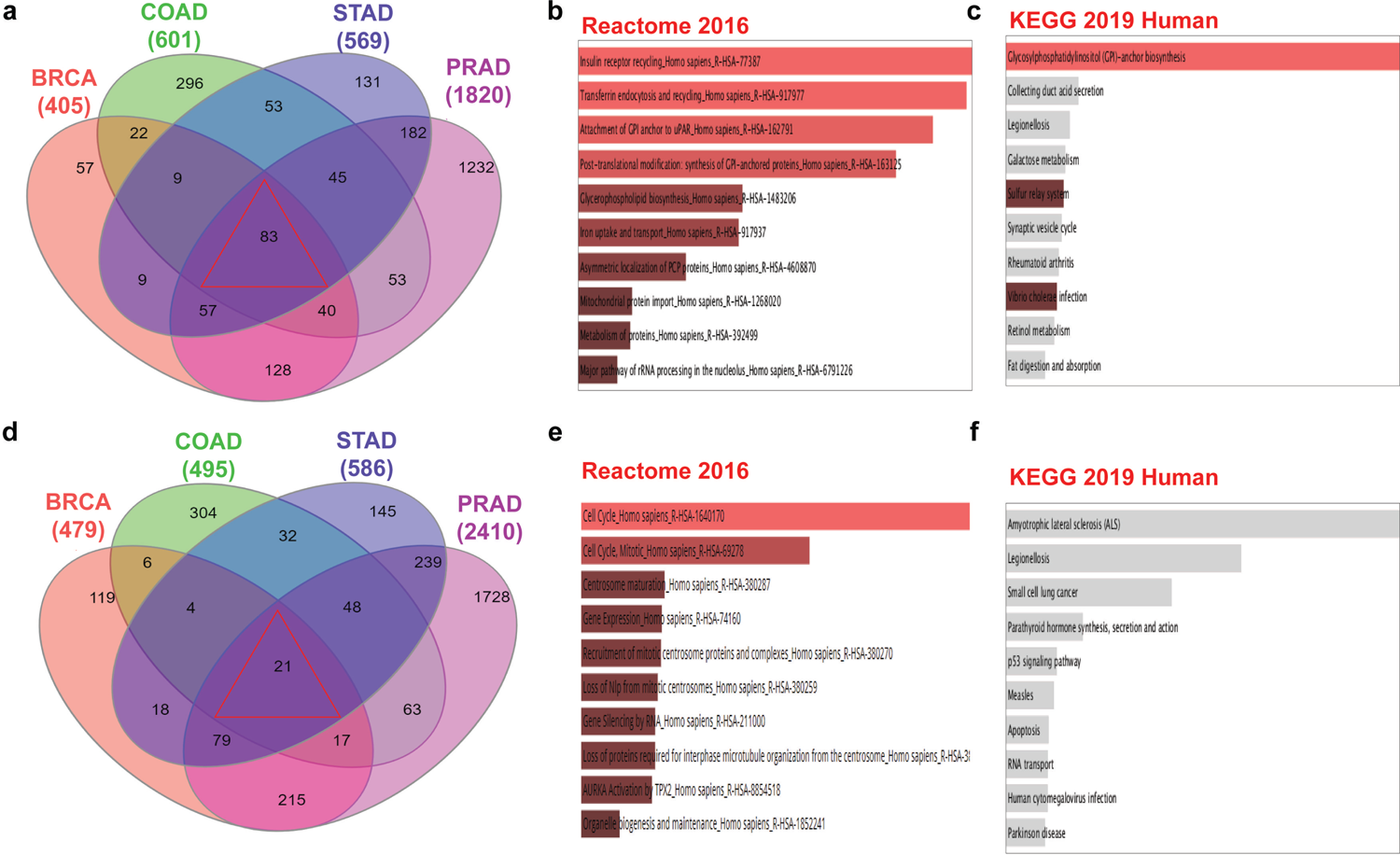
Pathway enrichment analysis for genes positively and negatively co-expressed with GRINA in breast, colon, stomach, and prostate cancer. (**a**) Venn diagram of genes positively co-expressed with GRINA based on R2: Genomics Analysis and Visualization Platform databases for breast, colon, stomach, and prostate cancer, using InteractiVenn. (**b and c**) Pathway (Reactome and KEGG) enrichment analysis was performed using Enrichr for the 83 genes positively co-expressed with GRINA in breast, colon, stomach, and prostate cancer. (**d**) The Venn diagram of genes negatively co-expressed with GRINA based on R2: Genomics Analysis and Visualization Platform databases for breast, colon, stomach, and prostate cancer using InteractiVenn web. (**e and f**) Pathway (KEGG and Reactome) enrichment analysis was performed using Enrichr web for the 21 genes negatively co-expressed with GRINA in breast, colon, stomach, and prostate cancer.

We performed Reactome and KEGG (Kyoto Encyclopedia of Genes and Genomes) pathway analysis for both the positive and negative clusters. Our Reactome pathway analysis revealed that some positive cluster genes were associated with pathways related to cell surface events, including insulin receptor recycling, transferrin endocytosis and recycling, and attachment of GPI anchors to uPAR (Figure 8b). Some categories involved post-transcriptional protein regulation, such as the synthesis of GPI-anchored proteins. For negative cluster genes, pathways related to the cell cycle and transcriptional regulation, such as cell cycle, gene expression, and gene silencing by RNA, predominated (Figure 8e). KEGG analysis indicated that glycosylphosphatidylinositol (GPI)-anchor biosynthesis, which was also found in Reactome pathway analysis, was the prominent pathway for the positive cluster (Figure 8c). In addition, KEGG analysis indicated that some negative cluster genes were associated with p53 signaling, apoptosis, and RNA transport (Figure 8f). Thus, our findings suggested that the biological pathways involved in regulating genes positively co-expressed with GRINA are fundamentally different from those showing negative co-expression with GRINA. GRINA may be associated with specific critical pathways related to cell surface dynamics and post-transcriptional control in cancer development.

### 3.9. Upstream Regulator Analysis of Genes Co-expressed with GRINA

We also identified transcription factors, intermediate proteins, and associated protein kinases potentially related to regulating co-expression of genes with *GRINA* using the Expression2Kinases (X2K) platform. X2K analysis revealed that PML, HNF4A, and TAF1 are major transcription factors that bind to positive cluster genes (Figure 9a; Supplementary Table 3), whereas STAT3, PPARG, and EGR1 are prominent transcription factors associated with negative cluster genes (Figure 9b; Supplementary Table 3). Transcription factors associated with upregulated genes were generally distinct from those associated with downregulated genes except for TAF1.

**Figure 9.**
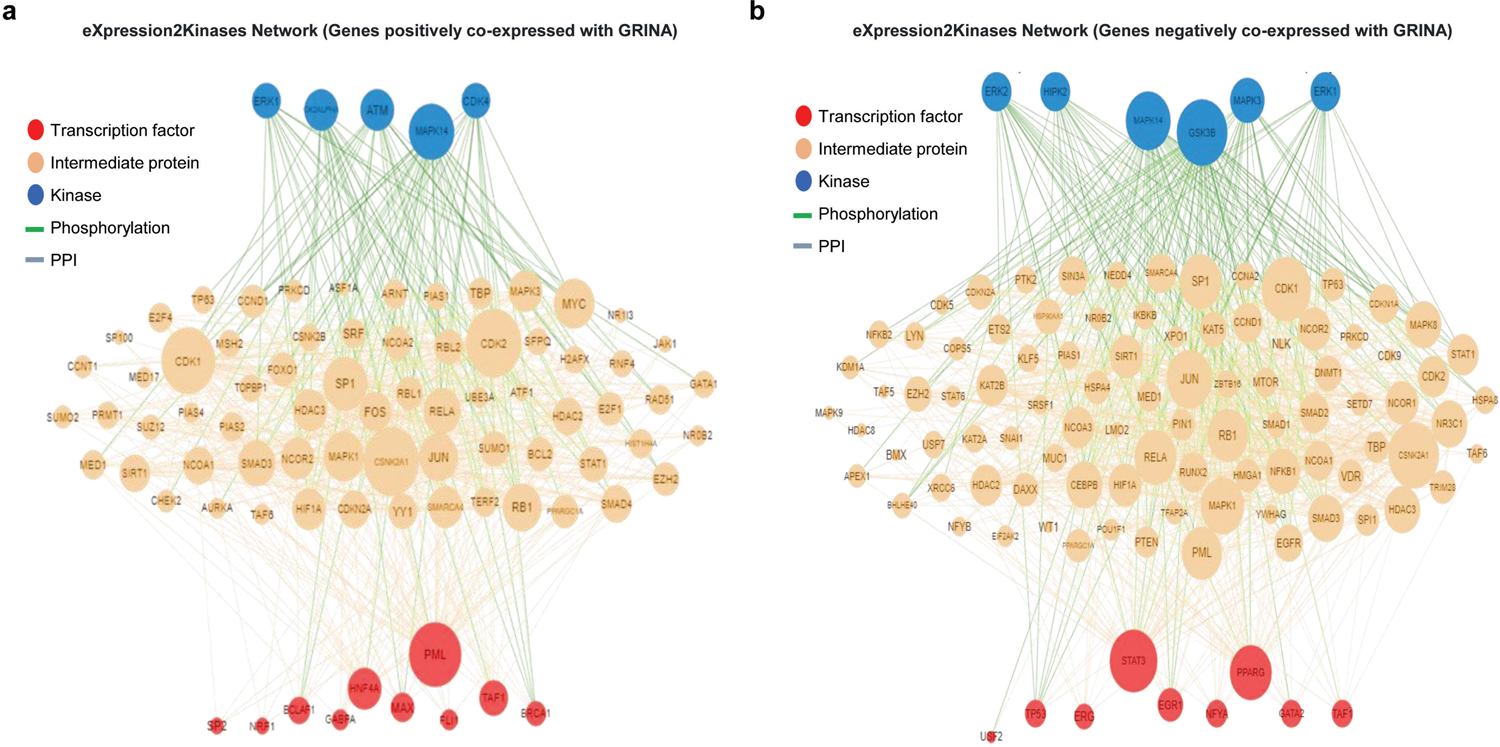
Subnetwork of transcription factors, intermediate proteins, and protein kinases. Expression2Kinases analysis of the (**a**) positively- and (**b**) negatively co-expressed genes indicated the most enriched TFs and kinase upstream of co-expressed genes occurring in multiple cancers based on a combination of P-values and *z*-scores. Node size reflects connectivity, and color indicates transcription factors in red, intermediate proteins in orange, and kinases in blue.

We also noticed that these transcription factors of interest for both the positive and negative clusters related to a large number of intermediate proteins that may assist with transcription factor activation. In addition, protein kinases such as MAPK14, ATM, and CK2ALPHA for positively co-expressed genes (Figure 9a; Supplementary Table 4) and GK3B, MAPK14, and MAPK3 for negatively co-expressed genes (Figure 9b; Supplementary Table 4) showed connections with many intermediate proteins associated with transcription factors belonging to each cluster. The only protein kinase in common between the positive and negative clusters was MAPK14. Promyelocytic leukemia protein (PML) and signal transducer and activator of transcription 3 (STAT3) showed the largest numbers of direct/indirect connections. They were labeled hub proteins of the positive and negative cluster genes, respectively. Our findings suggest that candidate hub proteins and their downstream targets may play significant roles in the progression of breast, colon, gastric, and prostate cancers. These transcription factors thus can be considered potential biomarkers and may be utilized in targeted therapy for the above cancers.

## 4. Discussion

*GRINA* belongs to the TMBIM protein family and is involved in calcium homeostasis, regulating various vital processes such as cell survival and neurotransmitter release. Alterations in the roles of *GRINA* are associated with schizophrenia and celiac disease [41]. Irregular *GRINA* expression patterns are found in several cancers, and elevated *GRINA* levels promote gastric cancer growth [13]. In contrast, *GRINA* is downregulated in the post-mortem superior temporal gyrus of schizophrenia patients [46]. Although its relevance is still unknown, alternative splicing of *GRINA* is found in the cortex of Alzheimer’s patients and cutaneous horn cancer [47]. GRINA is one of the most common NMDA receptors. NMDA receptors’ expressions were reported in human ovarian cancer tissues and human ovarian cancer cell lines [8]. As they were expressed in breast cancer and small-cell lung cancer, these receptors can potentially be targeted to trigger cancer cell death [3, 4]. Overexcitation of the extrasynaptic receptors in the peritumoral neurons was linked to the development of peritumoral seizures [48]. GRINA is also predominantly expressed in the brain [49]. In our current study, we analyzed *GRINA* expression in various cancers using bioinformatics tools and multiple gene expression datasets. Our analysis showed that *GRINA* expression is significantly elevated in multiple cancer cells compared to their normal counterparts. The consistency of *GRINA* expression patterns in each cancer of interest was crosschecked using different databases and confirmed based on *GRINA* promoter and immunostaining results. *GRINA* mRNA levels positively correlated with alteration frequency and negatively correlated with patient survival in breast, colon, gastric, and prostate cancers. These results suggest that *GRINA* function significantly affects the prognosis and progression of various cancers. In addition, the consistent elevation of *GRINA* expression in cancers indicates some commonality amongst different cancer types where *GRINA*-centric regulatory mechanisms may function. As *GRINA* expression is variable and has a wide range in many tumors, the question then comes to interpreting this variability in the clinical setting. In molecular therapeutic target viewpoint, the elevated expression of *GRINA* in the cancer of interest first conveys biomarker significance. Because of the confirmed augmented expression, the physician might be more confident to apply the targeted therapies. Then, the strength to which these targeted drugs should be clinically applied does partly come from GRINA expression variability. However, adequate prior clinical trials must be conducted to see whether they can be approved for broader use. The existence of co-expressed genes with *GRINA* can appear as additional confirmation of the biomarker role of *GRINA* and the strength of the *GRINA*-mediated targeted therapies.

To reveal potential *GRINA*-related molecular mechanisms and signaling pathways, we examined *GRINA* miRNA-mRNA regulatory networks, the identities and function of genes co-expressed with *GRINA*, and upstream regulator analysis of those co-expressed genes. Analysis of miRNA-mediated post-transcriptional regulation of *GRINA* mRNAs identified miR-411-5p, miR-654-5p, and miR-874-3p as miRNAs of interest, the expression of which was significantly lower in breast and prostate cancers, following the ceRNA model. Relatively low expression levels of these miRNAs were also noticed in the colon and gastric, with some minor exceptions. There exist some evidence that miRNA post-transcriptionally regulates *GRINA* expression at both mRNA and protein levels. A study reported that several microRNAs such as miR-411 were predicted to target alcohol-responsive mRNAs, including *GRINA* [50].

Interestingly, miR-411 is also found in our miRNA-mRNA analysis. *GRINA* was also significantly upregulated in human trabecular meshwork cells after overexpression of miR-24 mimic [51]. Moreover, *GRINA* has an important activity controlling cell death induced by ER stress suggesting a functional interconnection between *GRINA* and its role in the control of cell death and ER calcium homeostasis [52]. Our analysis indicates that a regulatory network consisting of miRNAs, including miR-411-5p, miR-654-5p, and miR-874-3p, may facilitate the oncogenic roles of *GRINA* in certain cancers.

Co-expression analysis may also provide important information for investigating mechanisms of *GRINA* function. A positive correlation between the expression of several genes and *GRINA* levels was observed in breast, colon, gastric, and prostate cancers, 83 of which occurred in all cancers. In contrast, the number of negatively co-expressed genes present in all cancers examined was 21. Functional enrichment analysis of positively co-expressed genes revealed that GPI anchor to protein, GPI anchor binding, and mRNA cleavage and polyadenylation specificity factor complex were the most significantly enriched GO terms. Reactome pathway analysis showed that insulin receptor recycling, transferrin endocytosis and recycling, attachment of GPI anchor to uPAR, and post-translational modification and synthesis of GPI-anchored proteins were the major pathways associated with the positively co-expressed genes. Cell surface dynamics and post-transcriptional modification of *GRINA* have also been previously reported [41, 53, 54]. Noticeably, GO terms and pathways related to genes negatively co-expressed with *GRINA* were completely different from those related to positively co-expressed genes. Our findings suggest that pathways involving cell surface dynamics and post-transcriptional regulation may underpin the role of *GRINA* in cancer.

We next performed upstream regulator analysis of genes co-expressed with *GRINA* to identify associated major transcription factors. This analysis revealed that transcription factors regulating positive and negative cluster genes were influenced by many intermediate proteins modulated by specific kinases. The major transcription factors binding to positive cluster genes were PML, HNF4A, and TAF1, while major transcription factors associated with negative cluster genes were STAT3, PPARG, and EGR1. In addition, PML and STAT3 acted as hub proteins of the positively correlated genes and negatively correlated genes, respectively. The roles of PML and STAT3 as hub proteins in the said cancers are also evident from previous research. For example, the study showed that PML promotes metastasis of TNBC [55], and targeting PML elicits its growth suppression [56]. Also, the expression level of PML is the potential to predict prostate cancer progression [57] and gastric cancer [58]. Furthermore, histochemical analyses of clinical samples have shown PML to be downregulated in colon cancer [59]. On the other hand, STAT3 might serve as a therapeutic target for various cancers such as gastric [60] and TNBC [61]. Our findings thus indicate that the PML could be used as a biomarker in *GRINA*-mediated targeted therapy for breast, colon, gastric, and prostate cancers.

In this study, we investigated the expression, methylation, genetic alteration, and immunostaining patterns of *GRINA*. We evaluated its prognostic use through organized data analysis using several established bioinformatics tools, with expression and clinical data obtained from various open-source platforms. Our study showed that the expression, methylation, and immunostaining of *GRINA* collectively convey biomarker significance for breast, colon, gastric, and prostate cancers, among others. The overexpression of *GRINA* was in agreement with its promoter methylation level in the aforementioned cancers. However, the results of hypomethylation of the *GRINA* gene promoter in breast and prostate cancers were more significant compared to that in colon and gastric cancers. Our analysis also indicates that high *GRINA* expression positively correlates with poor prognosis in patients with the said cancers. The silico data mining of miRNA reveals that the regulatory network containing miR-411-5p, miR-654-5p, and miR-874-3p may contributes to the oncogenic roles of *GRINA* to some extent. In sum, this study tried to gain some insights into the oncogenic roles of *GRINA* by revealing the respective miRNA-mRNA interactions, genes co-expressed with *GRINA* and their associated pathways, and specific kinases, transcription factors, and accompanying intermediate proteins that could collectively underpin the regulation of *GRINA* expression. However, it is worth to note that data mining only constitutes the first step of a scientific investigation. Since there is no prominent and useful study and review on GRINA with data analysis perspective, we thus became motivated to investigate the expression and related molecular mechanisms of the GRINA gene and find their relevance to prognostic significance through systematic data analysis using publicly available expression and clinical outcomes data. On that, further research is required to validate the outcomes of this research.

## 5. Conclusions

In our multiomics analysis of *GRINA* expression in cancer databases, we provide evidence of a relationship between altered *GRINA* expression and clinical outcomes. Our study reveals the significance of *GRINA* expression and possible *GRINA*-related molecular mechanisms and pathways in cancer progression. The findings of this study thus may offer valuable insights into the use of *GRINA* as a prospective therapeutic target for various cancers.

### Abbreviations

BLCA, bladder urothelial carcinoma; BRCA, invasive breast carcinoma; CESC, Cervical squamous cell carcinoma; CHOL, cholangiocarcinoma; COAD, colon adenocarcinoma; ESCA, esophageal carcinoma; GBM, glioblastoma multiforme; HNSC, head and neck squamous cell carcinoma; KICH, kidney chromophobe; KIRC, kidney renal clear cell carcinoma; KIRP, Kidney renal papillary cell carcinoma; LIHC, Liver hepatocellular carcinoma; LUAD, Lung adenocarcinoma; LUSC, Lung squamous cell carcinoma; PAAD, pancreatic adenocarcinoma; PRAD, Prostate adenocarcinoma; PCPG, Pheochromocytoma and Paraganglioma; READ, rectum adenocarcinoma; SARC, Sarcoma; SKCM, Skin Cutaneous Melanoma; THCA, thyroid carcinoma; THYM, Thymoma; STAD, stomach adenocarcinoma; UCEC, uterine corpus endometrial carcinoma; LAML, Acute Myeloid Leukemia; OV, Ovarian serous cystadenocarcinoma; LBC, lobular breast cancer; IDBC, invasive ductal breast cancer; HER2, human epidermal growth factor receptor 2; TNBC, triple-negative breast cancer; CC, colon cancer; RADC, rectal adenocarcinomas; COAD, colon adenocarcinoma; MADC, mucinous-adenocarcinoma; GM, Gastric Mucosa; GITA, Gastric Intestinal Type Adenocarcinoma; GMA, Gastric Mixed Adenocarcinoma; STAD, stomach adenocarcinoma; PG, Prostate Gland; PC, Prostate Carcinoma; PIN, Prostatic Intraepithelial Neoplasia; GS, Gleason scores.

### Author Contributions

Conceptualization, S.K.S.; methodology, S.K.S., and S.M.R.I.; validation, S.K.S., and S.M.R.I.; formal analysis, S.K.S., S.M.R.I., S.E.-S., F.T., J.D., M.A., and S.-G.C; investigation, S.K.S., and S.M.R.I.; data curation, S.K.S., and S.M.R.I.; Funding acquisition, S.-G.C.; writing—original draft preparation, S.M.R.I.; writing—review and editing, S.K.S.; visualization, S.K.S.; supervision, S.K.S., and S.M.R.I. All authors have read and agreed to the published version of the manuscript.

## Acknowledgments

We are grateful to the contributors of data to Oncomine, UCSC Xena, ULCAN, Human Protein Atlas, cBioPortal, R2: Kaplan Meier Scanner web, SurvExpress web, miRSystem, starBase v3.0, InteractiVenn, Enrichr web, and Expression2Kinases (X2K) web, which provides a Web resource for exploring, visualizing, and analyzing multidimensional cancer genomics data.

## Supplementary Information

**Supplementary Figure 1.**
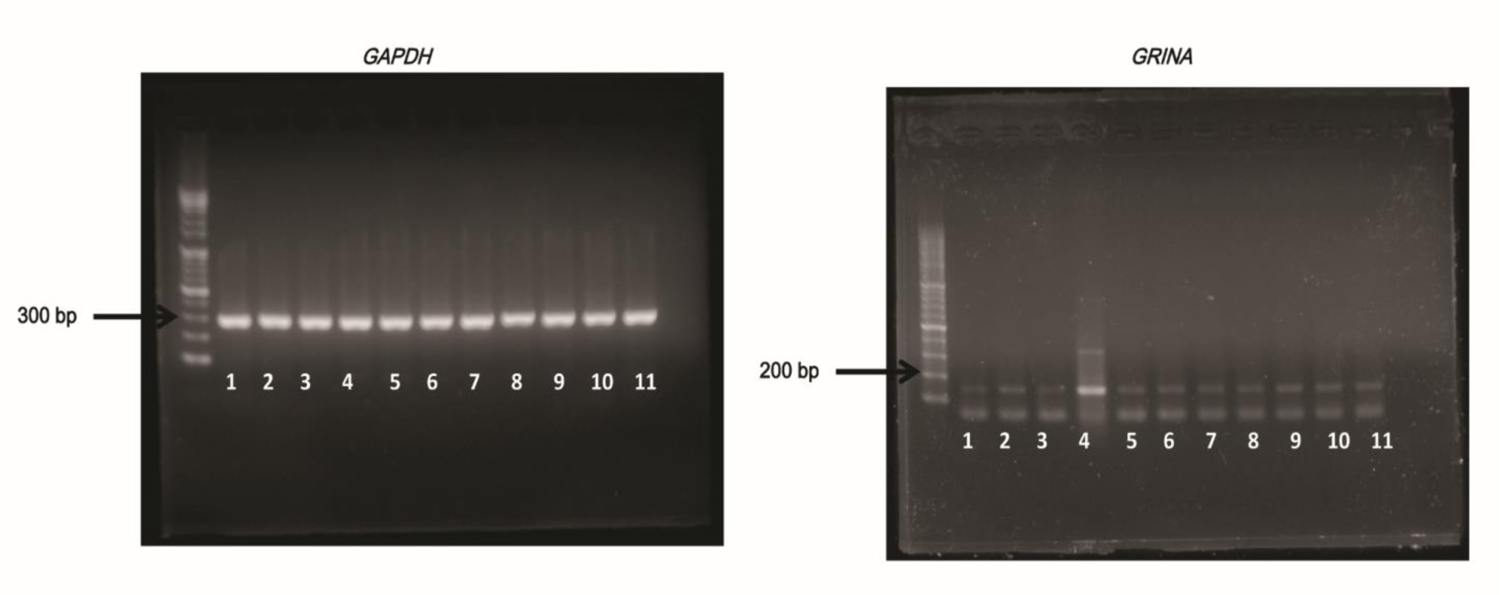
Agarose gel picture of GRINA and GAPDH. Numbers indicate the cell lines added in Figure 2. Arrows indicate the size of PCR product.

**Supplementary Figure 2.**
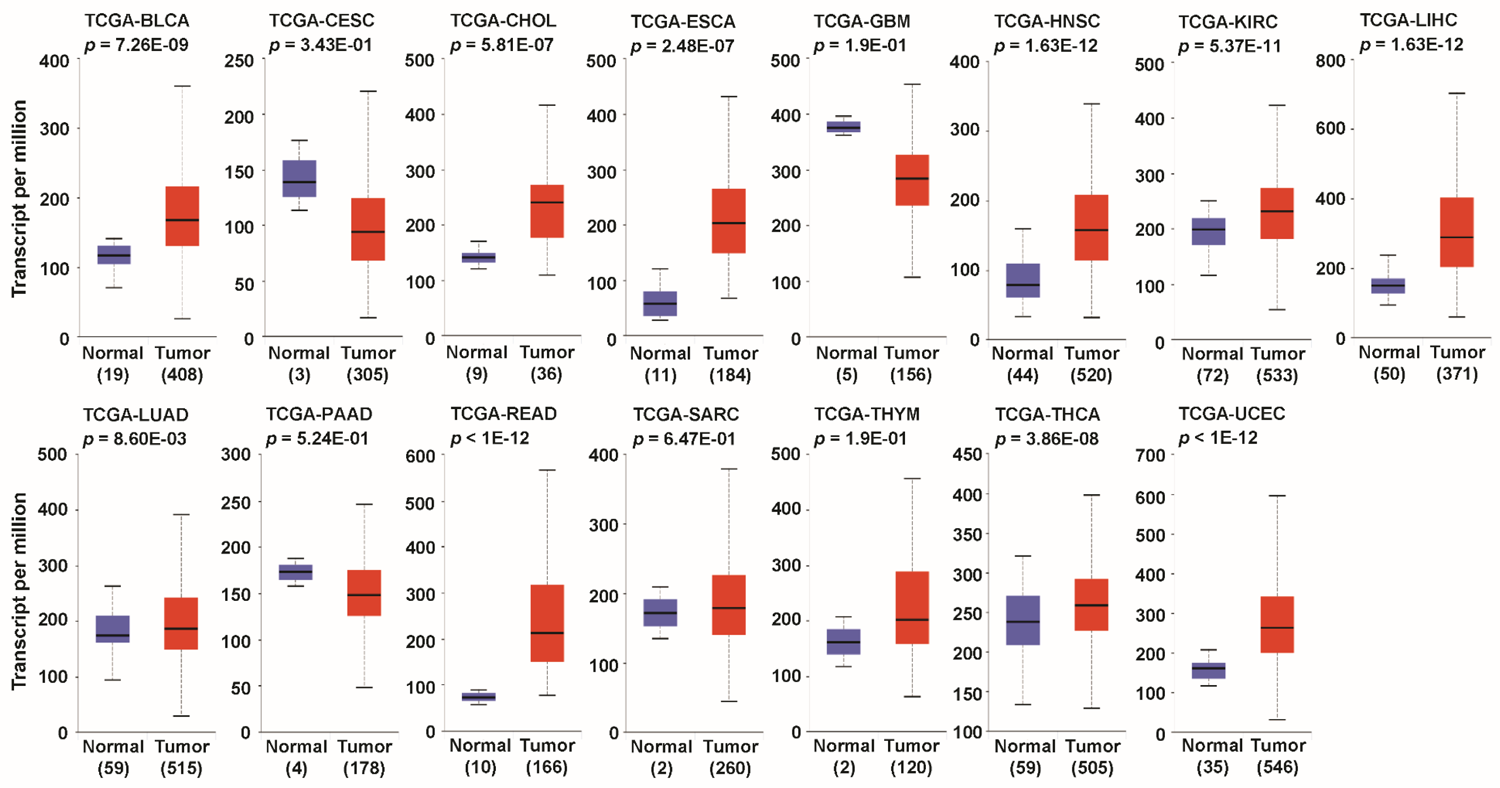
*GRINA* mRNA expression analysis in different cancer types (TCGA database). *GRINA* expression levels from data in the Cancer Genome Atlas (TCGA) database. Box plots showing *GRINA* mRNA levels in various tumours and corresponding normal tissues derived from data in TCGA database through ULCAN. (Abbreviations: BLCA-bladder urothelial carcinoma; CESC-cervical squamous cell carcinoma and endocervical adenocarcinoma; CHOL-cholangial carcinoma; ESCA-oesophageal carcinoma; GBM-glioblastoma multiforme; HNSC-head and neck squamous cell carcinoma; KIRC-kidney renal clear cell carcinoma; LIHC-liver hepatocellular carcinoma; LUAD-lung adenocarcinoma; PAAD-pancreatic adenocarcinoma; READ-rectum adenocarcinoma; SARC-sarcoma; THYM-thymoma; THCA-thyroid carcinoma; UCEC-uterine corpus endometrial carcinoma).

**Supplementary Figure 3.**
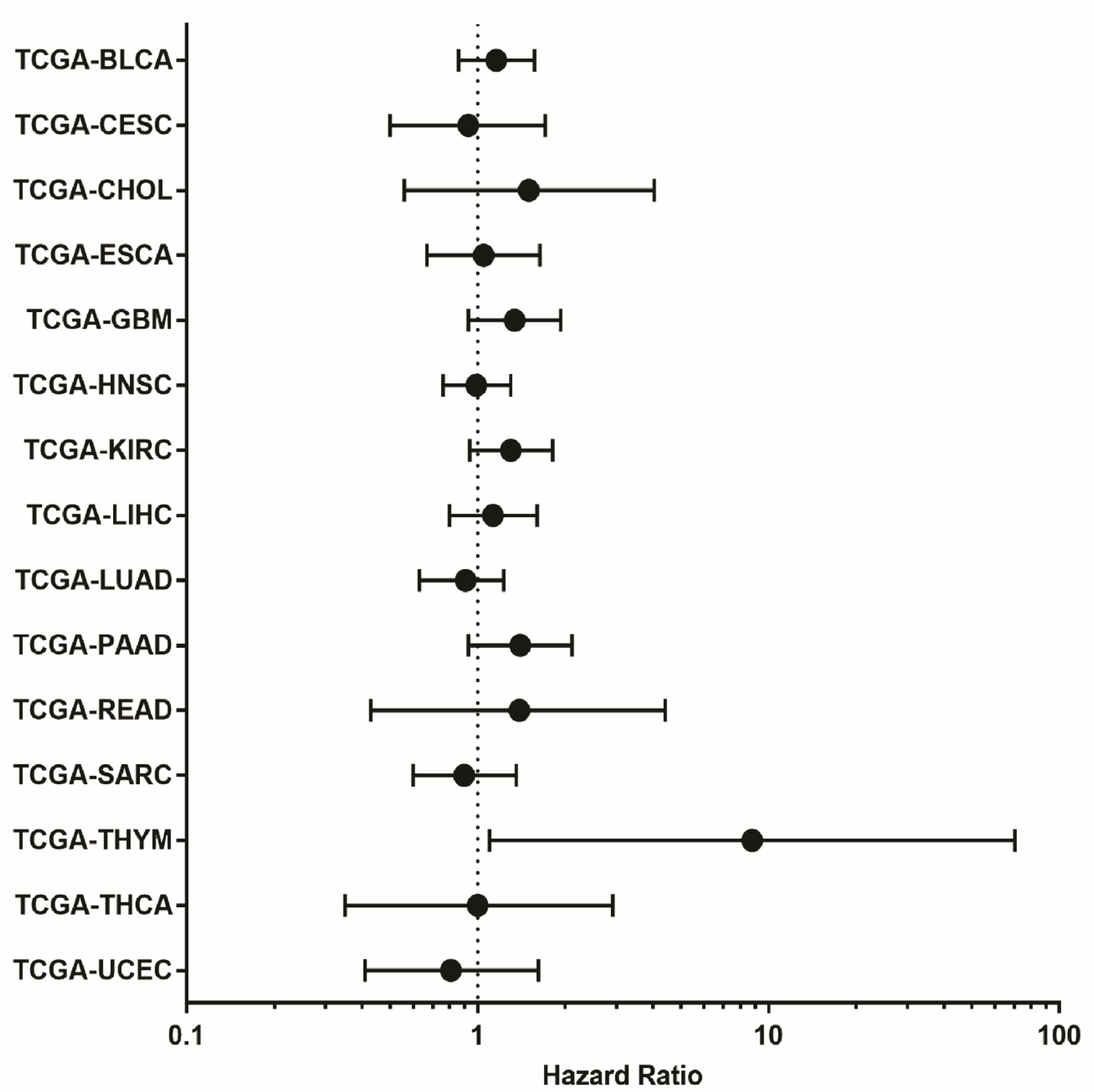
Prognostic relevance of GRINA mRNA levels in various cancers based on TCGA data and SurvExpress (Available at http://bioinformatica.mty.itesm.mx:8080/Biomatec/SurvivaX.jsp). Survival curve analysis with a threshold of cox P-value < 0.05.

**Supplementary Figure 4.**
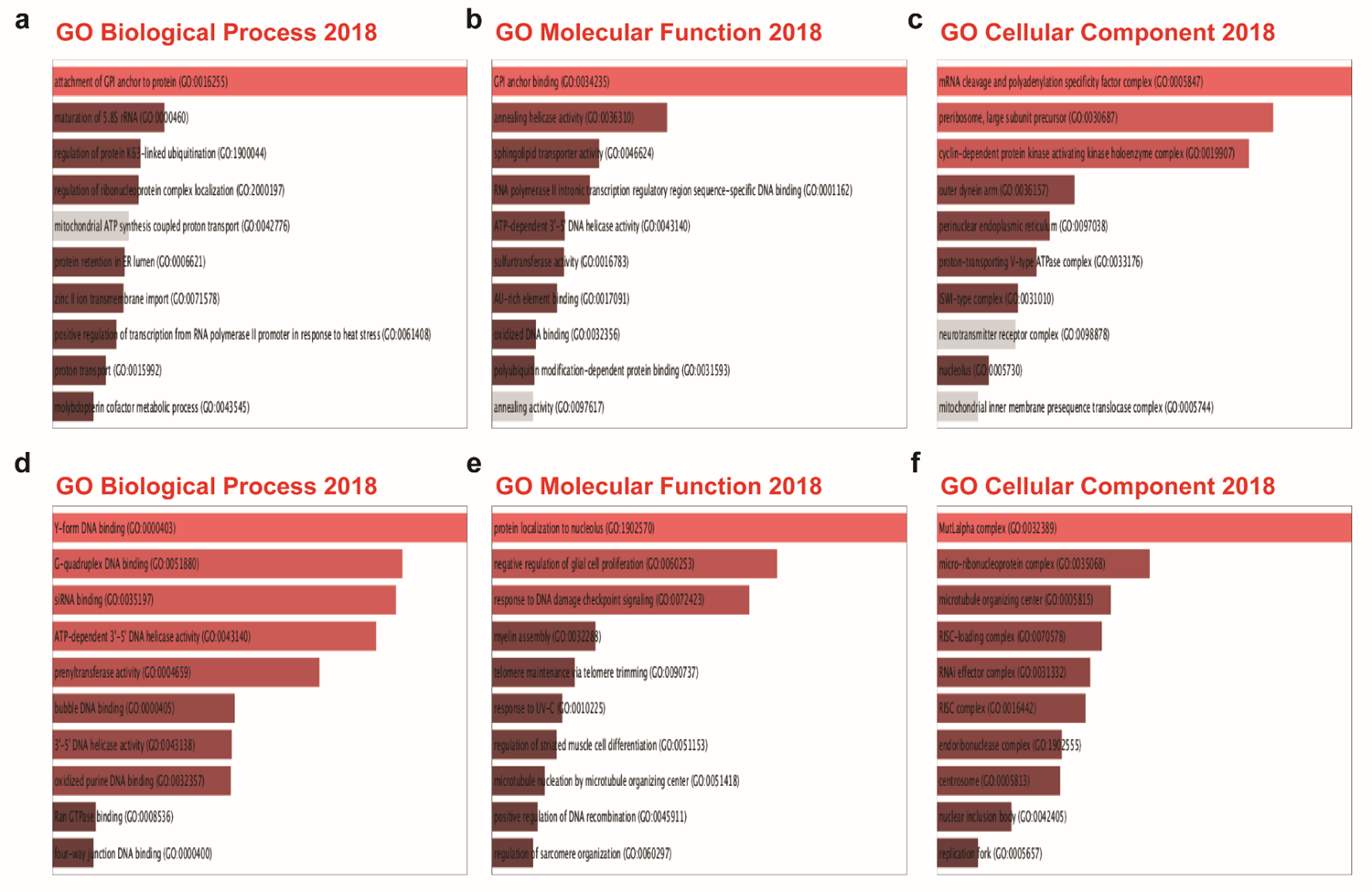
GO functional annotation analysis for genes positively and negatively co-expressed with GRINA in breast, colon, stomach, and prostate cancer. (**a-c**) GO functional annotation (biological process, molecular function, and cellular component) was performed using Enrichr for 83 genes positively co-expressed with GRINA in breast, colon, stomach, and prostate cancer. (**d-f**) GO functional annotation (biological process, molecular function, and cellular component) was performed using Enrichr web for the 21 genes negatively co-expressed with GRINA in breast, colon, stomach, and prostate cancer.

**Supplementary Table 1.**
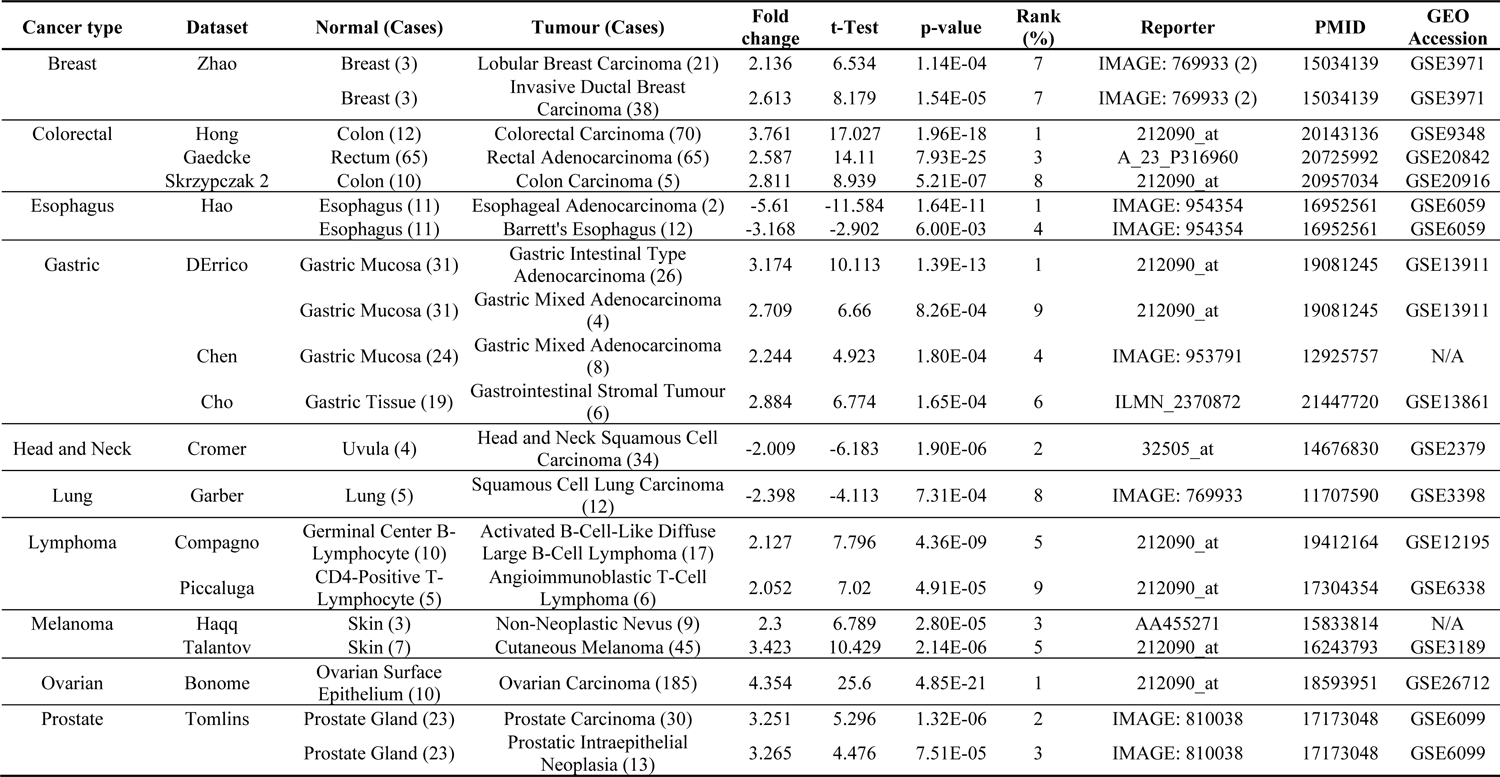
Changes in GRINA expression at the transcriptional level between different types of cancer and normal tissues (Oncomine Database).

**Supplementary Table 2.** Changes in GRINA expression at the transcriptional level between different types of cancer and normal tissues (TCGA Database).

| Order | Cancer type | # of normal samples | # of cancer samples | GRINA expression in cancer (High/Low) | p-value (Welch's t-test) |
| --- | --- | --- | --- | --- | --- |
| 1 | Bladder urothelial carcinoma (BLCA) | 19 | 408 | High | 5.4E-06 |
| 2 | Breast invasive carcinoma (BRCA) | 114 | 1097 | High | 4.6E-38 |
| 3 | Cervical squamous cell carcinoma (CESC) | 3 | 305 | Low | 0.3771 |
| 4 | Cholangiocarcinoma (CHOL) | 9 | 36 | Low | 3.05E-06 |
| 5 | Colon adenocarcinoma (COAD) | 4 | 286 | High | 2.25E-22 |
| 6 | Esophageal carcinoma (ESCA) | 11 | 184 | High | 4.81E-06 |
| 7 | Glioblastoma multiforme (GBM) | 5 | 156 | Low | 0.289002 |
| 8 | Head and neck squamous cell carcinoma (HNSC) | 44 | 520 | High | 1.52E-10 |
| 9 | Kidney chromophobe (KICH) | 25 | 67 | High | 2.42E-09 |
| 8 | Kidney renal clear cell carcinoma (KIRC) | 72 | 533 | High | 1.28E-12 |
| 11 | Kidney renal papillary cell carcinoma (KIRP) | 32 | 290 | Low | 0.386717 |
| 12 | Liver hepatocellular carcinoma (LIHC) | 50 | 371 | High | 4.65E-10 |
| 13 | Lung adenocarcinoma (LUAD) | 59 | 515 | High | 0.278277 |
| 14 | Lung squamous cell carcinoma (LUSC) | 52 | 503 | Low | 0.014681 |
| 15 | Pancreatic adenocarcinoma (PAAD) | 4 | 178 | Low | 0.432492 |
| 16 | Prostate adenocarcinoma (PRAD) | 52 | 497 | High | 5.12E-10 |
| 17 | Pheochromocytoma and Paraganglioma (PCPG) | 3 | 179 | Low | 0.240684 |
| 18 | Rectum adenocarcinoma (READ) | 10 | 166 | High | 7.18E-14 |
| **19** | Sarcoma (SARC) | 2 | 260 | Low | N/A |
| **20** | Skin Cutaneous Melanoma (SKCM) | 1 | 472 | Low | N/A |
| **21** | Thyroid carcinoma (THCA) | 59 | 505 | High | 9.3E-15 |
| **22** | Thymoma (THYM) | 2 | 120 | High | N/A |
| **23** | Stomach adenocarcinoma (STAD) | 34 | 415 | High | 7E-09 |
| **24** | Uterine corpus endometrial carcinoma (UCEC) | 35 | 546 | High | 6.1E-12 |

**Supplementary Table 3.**
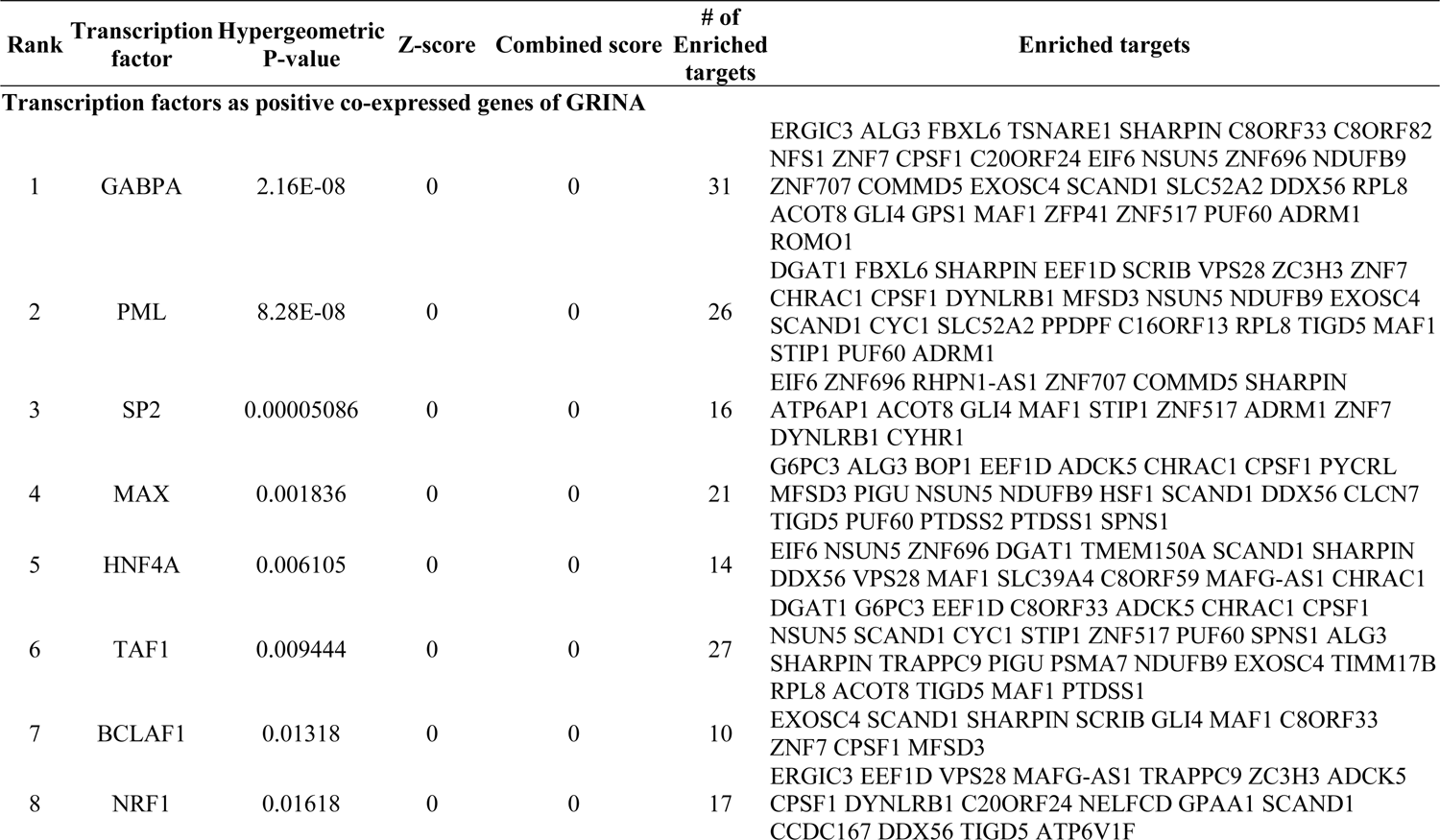
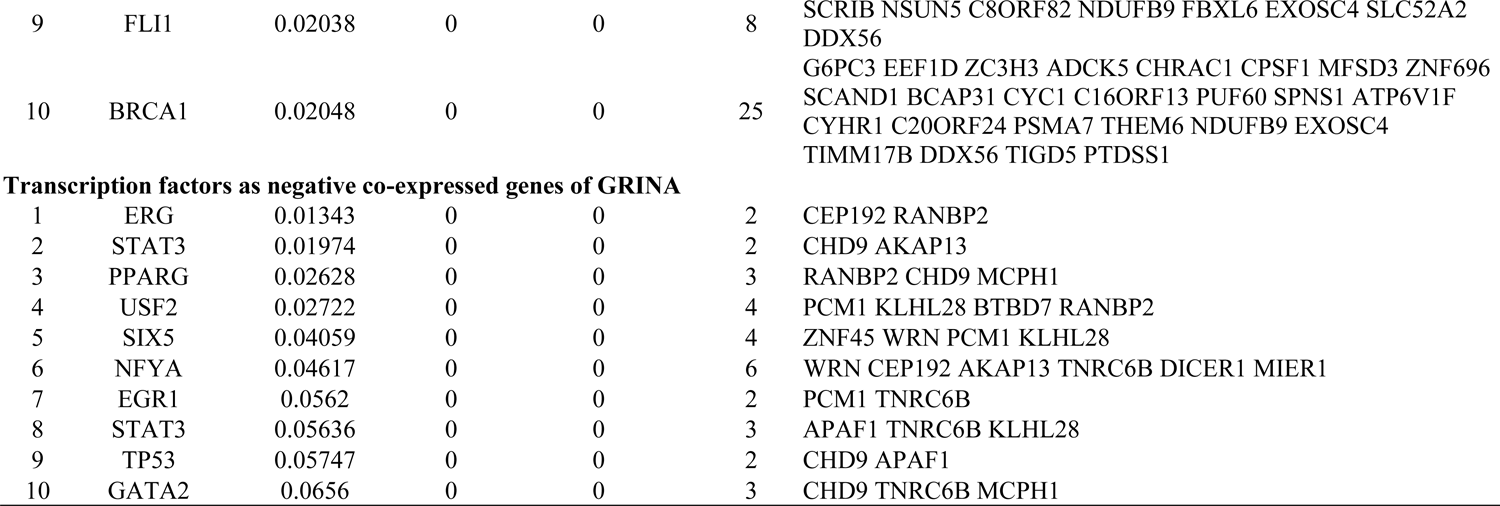
Transcription factors regulating genes positively and negatively co-expressed with GRINA identified using Expression2Kinases (X2K).

**Supplementary Table 4.**
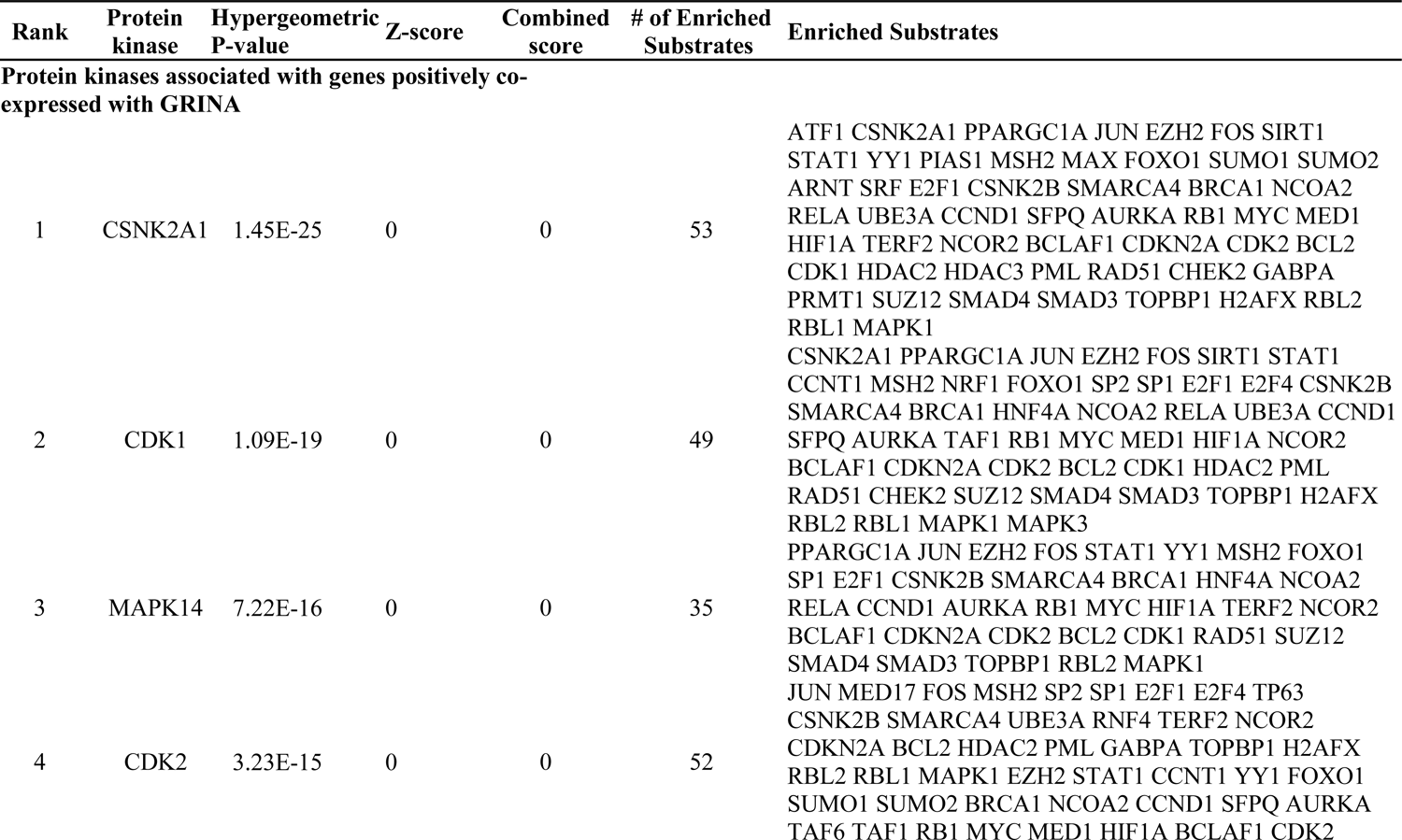
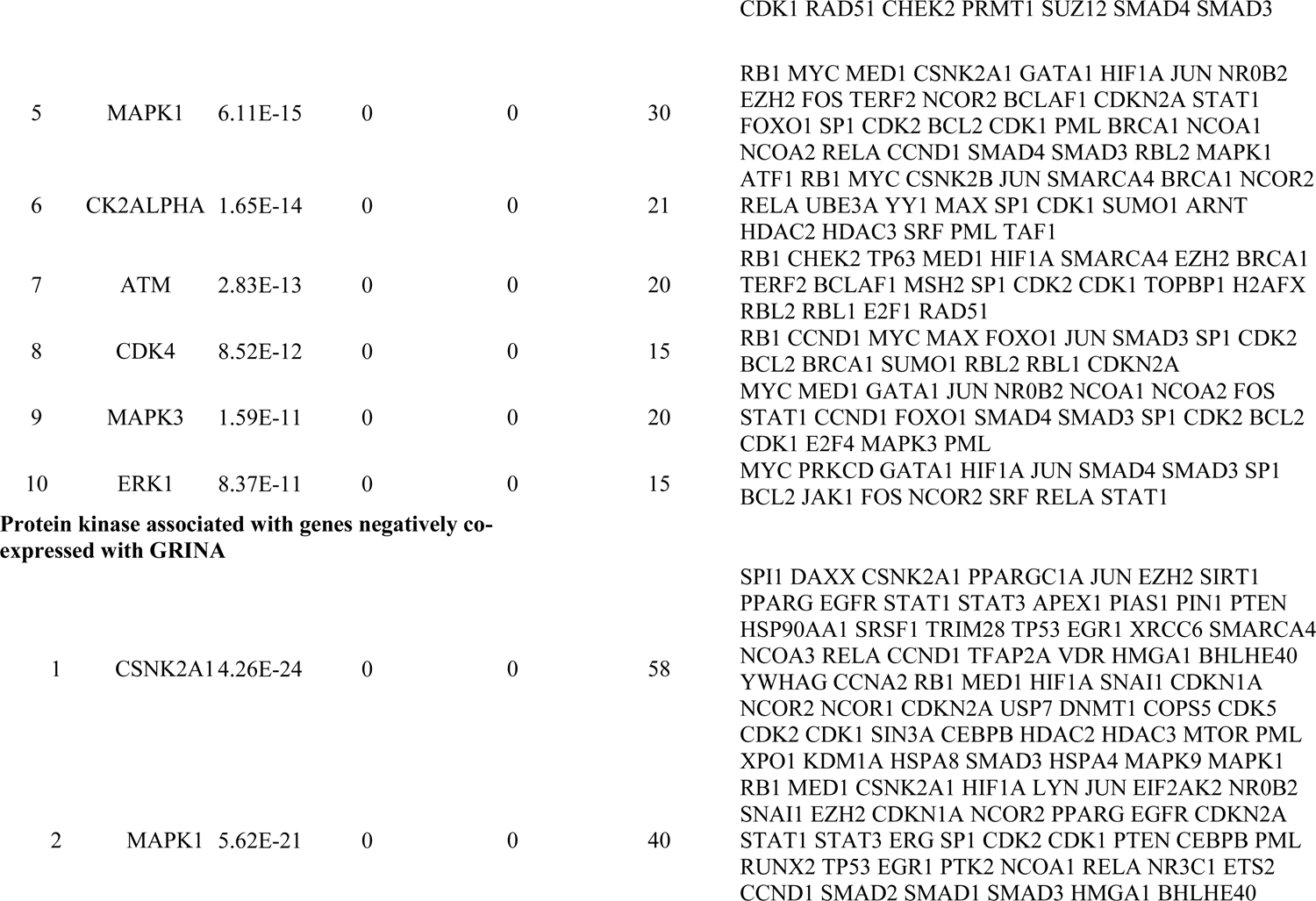
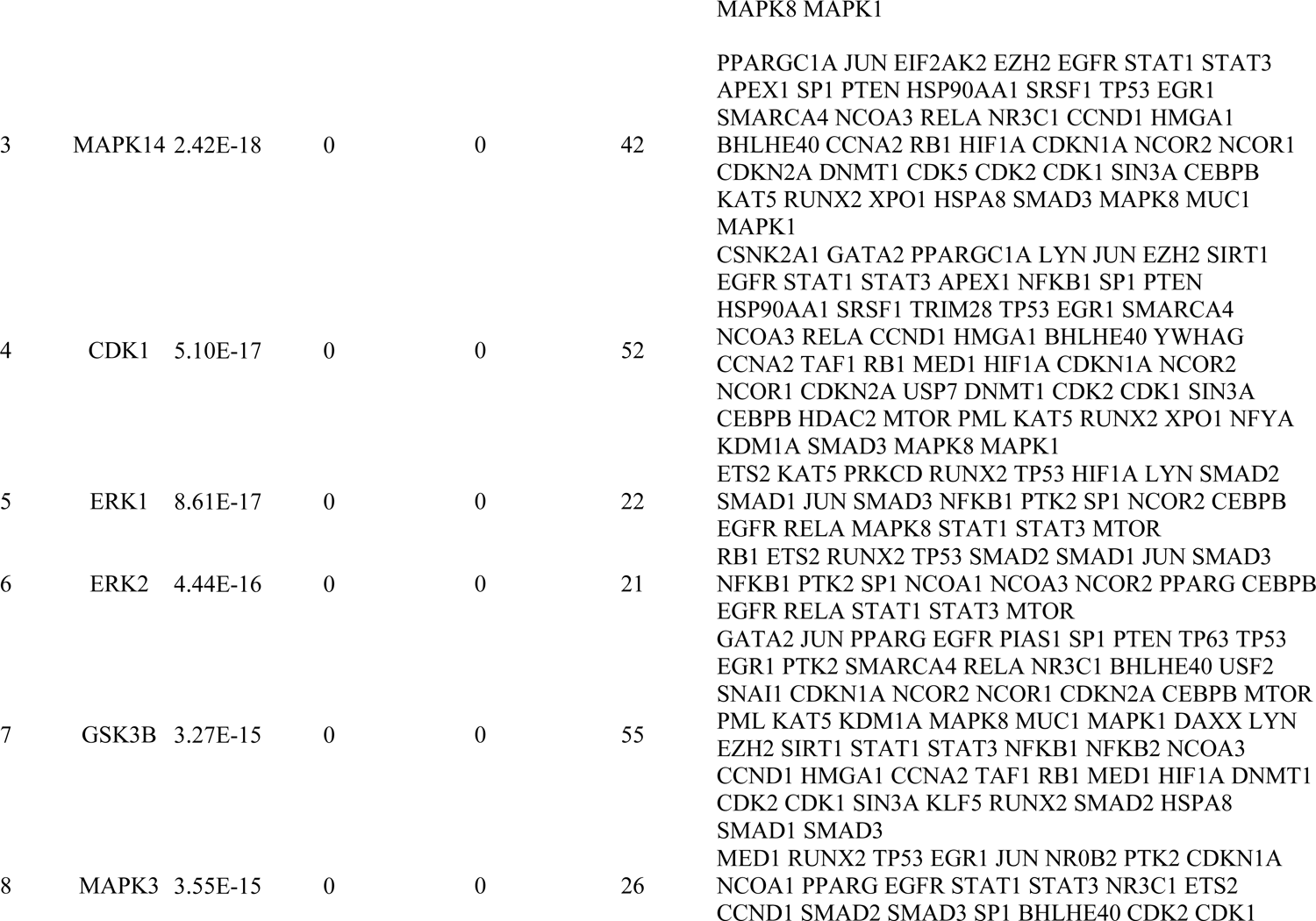
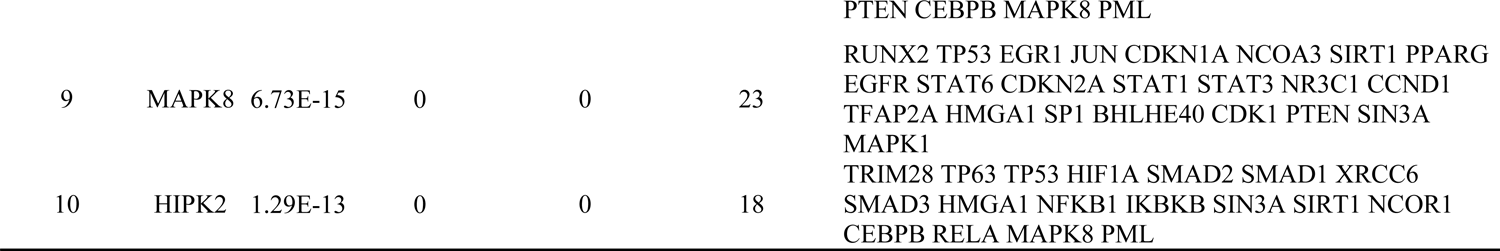
Protein kinases likely acting on genes positively- and negatively co-expressed with GRINA that are responsible for phosphorylation of PPI in various cancers.

